# StaVia: Spatially and temporally aware cartography with higher order random walks for cell atlases

**DOI:** 10.1101/2024.01.29.577871

**Authors:** Shobana V. Stassen, Minato Kobashi, Yuanhua Huang, Joshua W. K. Ho, Kevin K. Tsia

**Affiliations:** Department of Electrical & Electronic Engineering, The University of Hong Kong, Pokfulam Road, Hong Kong; School of Biomedical Sciences, Li Ka Shing Faculty of Medicine, The University of Hong Kong, Pokfulam, Hong Kong; Advanced Biomedical Instrumentation Centre, Hong Kong Science Park, Shatin, New Territories, Hong Kong; Laboratory of Data Discovery for Health, Hong Kong Science Park, Shatin, New Territories, Hong Kong; Department of Statistics and Actuarial Science, University of Hong Kong, Hong Kong

## Abstract

Single-cell atlases are critical for unraveling the cellular basis of health and disease, yet their sheer diversity and vast data landscape in time and space pose daunting computational challenges pertaining to the delineation of multiple simultaneously emerging lineages, the integration of spatial and temporal information, and the visualization of trajectories across large atlases with enough resolution to observe localized transitions. To tackle this intricacy, we introduce StaVia, a computational framework that synergizes multi-faceted single-cell data—spanning time-series data, spatial gene expression patterns, and directional trends from RNA velocity— with higher-order random walks that leverage the memory of cells’ past states. StaVia fuses this method with a cartographic “Atlas View” that offers intuitive graph visualization, simultaneously capturing the nuanced details of cellular development at single-cell resolution as well as the broader connectivity of cell lineages, avoiding common pitfalls of merged distinct trajectories or missed transitional states seen in existing methods which are all memoryless. Notably, we demonstrate that StaVia unlocks new insights into placode development, radial glia pluripotency during neurulation, and the transitions pivotal to these processes in a large-scale Zebrafish developmental atlas. StaVia also allows spatially aware cartography that captures relationships between cell populations based on their spatial location as well as their gene expressions in a MERFISH dataset - underscoring its potential to dissect complex biological landscapes in both spatial and temporal contexts.

## Introduction

The recent surge in the creation of single-cell atlases has ushered in a new era of understanding the complexities of life at the cellular level. These atlases are now instrumental for studying a wide range of tissues, organs, and even whole organisms to reveal the origins of cellular differentiation and functional diversity [Quake 2022, Tabula Sapiens Consortium 2022]. Further combined with spatial and time-series studies, they offer a high-definition window into biological development over space and time [Calderon 2022, Qiu 2023, Packer 2019, Sala 2019, Lange 2023]. However, the growing scale of single-cell atlases often poses daunting analytical challenges [Sikkema 2023, Qiu 2023, Lange 2023]. Specifically, the elevated complexity of large-scale atlases in terms of heterogeneity, temporal longitude, spatial environments, and sample sizes makes it difficult to unambiguously capture the emergence of multiple specialized cell lineages and their differentiation pathways at a high resolution, not to mention the difficulty of intuitively visualizing these complex pathways at this scale.

Available methods face three pressing challenges, the first is the inability to preserve localized details of underlying trajectories while maintaining a global view of their connectivity, resulting in differentiation pathways for distinct lineages being intermingled (by deviating into unrelated intermediate cell populations) or too myopic (failing to detect transition states). The second is that strategies to integrate available metadata (e.g spatial or temporal information) that could aid in the analysis of the cellular landscape are not readily available, thus forgoing the opportunity to use these sources of complementary information. Third, current practices to visualize developmental landscapes rely on established dimension-reduction visualization tools which primarily capture clusters of distinct lineages (e.g. UMAP [McInnes 2018], t-SNE [Maaten & Hinton 2008]). These tools are not designed to display a single-cell embedding that can intuitively be mapped or linked to inferred continuous trajectories. On the other hand, methods relying on diffusion maps (e.g. Phate [Moon 2019]) convey progression information at the expense of collapsing/superimposing multiple distinct lineages.

To address these challenges, we present StaVia, an automated end-to-end trajectory inference (TI) framework that uncovers cellular trajectories permeating large-scale single-cell spatial and temporal atlases without sacrificing the fine-grained details. To address the first obstacle, StaVia exploits a new form of lazy-teleporting random walks (LTRW) *with memory* to accurately pinpoint end-to-end trajectories in the atlas. Specifically, higher-order LTRW with memory are used to propagate information about a cell’s previous states when inferring subsequent states (e.g. during differentiation) (**Fig. 1a**). The inclusion of memory of past states critically alleviates issues seen in traditional first-order *memoryless* RW methods where pathways deviate into unrelated intermediate cell populations, or conversely become so localized that they fail to detect transition states **(Fig. 1b)**. Secondly, StaVia’s framework is also flexibly compatible with diverse input data types, in addition to RNA velocity, it offers seamless strategies to integrate spatial coordinates and temporal information (or other sequential metadata) to guide the cartography in a data driven manner **(Fig. 1a)**.

**Fig. 1.**
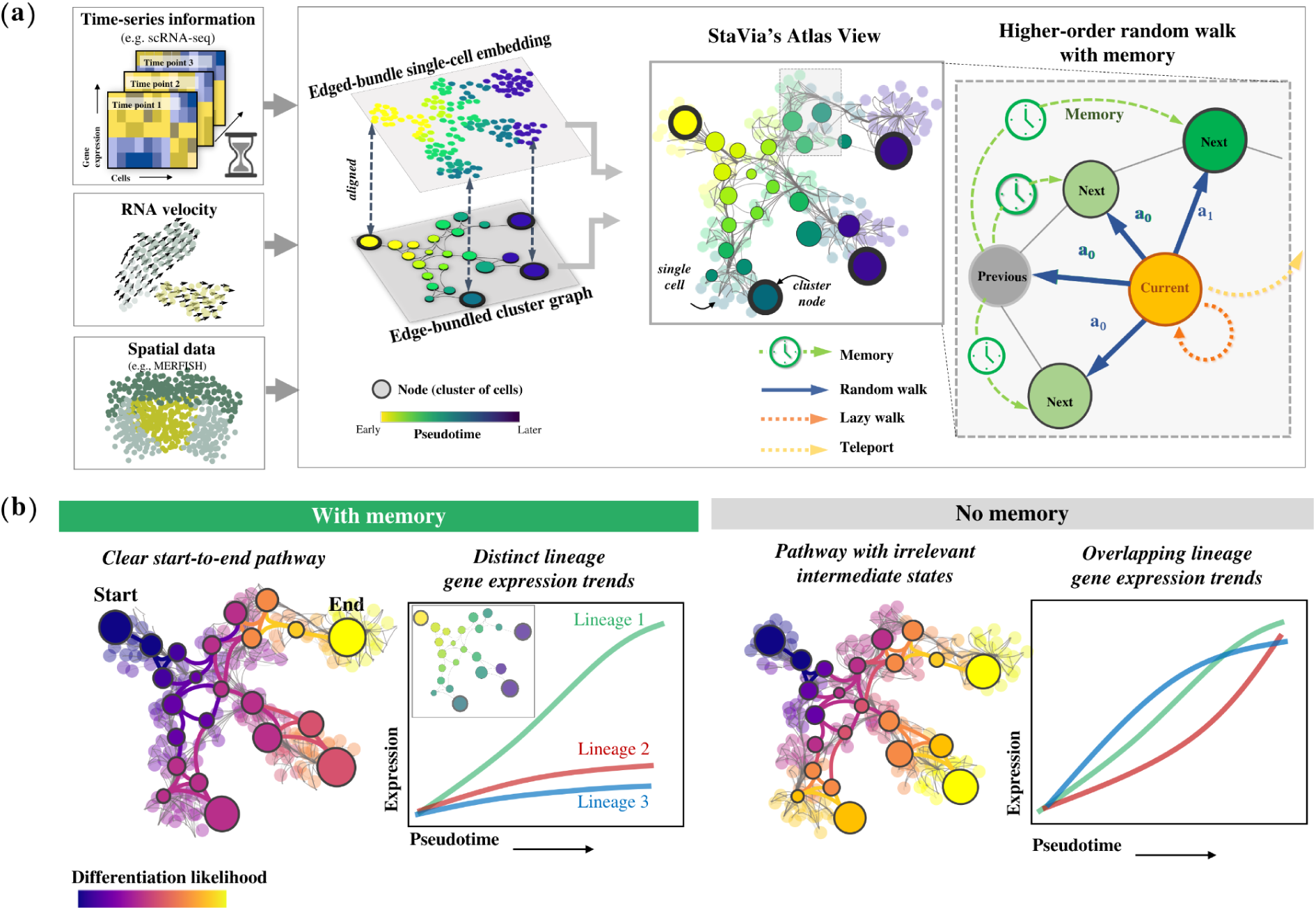
Overview of StaVia 2.0 Higher Order Random Walks with Memory. **(a)** The StaVia graph is a flexible framework for single-cell data that can optionally incorporate any combination of the following data to infer cell transitions: sequential or spatial metadata (e.g. known stages, tissue coordinates), RNA-velocity, pseudotime and lazy or teleporting behaviours. Based on an algorithm of higher order lazy-teleporting random walks (LTRW) with memory, StaVia can generate single-cell embeddings with the underlying high-resolution connectivity of the single-cell-KNN graph. This can be aligned with an edge-bundled cluster graph in which each node represents a cluster of single cells. The cluster graph and single-cell embedding can be overlaid to generate an Atlas View which offers an intuitive and comprehensive visualization for the computed trajectories. StaVia uses higher order lazy-teleporting random walks (LTRW) with memory to accurately infer complex trajectories. Previous states’ neighbors inform the decision making process for subsequent transitions, i.e. to determine the transition probabilities of moving from current state to the next states by introducing a memory factor **a** (**Methods**). **(b)** The Atlas View allows us to cartographically observe end-to-end pathways at a higher resolution. Higher-order LTRWs with memory ensure that pathways avoid detours to unrelated cell types and hence also increase the specificity of gene regulation along distinct lineages.

To address the third challenge of visually capturing complex TI landscapes, StaVia feeds forward the properties of the higher-order walks with memory and metadata to create a comprehensive cartographic *Atlas View*, which efficiently (both in terms of spatial layout and computational cost) integrates the high-resolution graph-edge information with the cell type specificity of single-cell embeddings to visually chart the predicted trajectories of entire atlases in a unified snapshot **(Fig. 1a)**. As a result, StaVia can simultaneously capture smooth sequential processes while maintaining the separation of distinct lineages - outperforming popular visualization tools (t-SNE, UMAP, Phate, etc.).

We use a murine gastrulation atlas [Pijuan-Sala 2019] and the recent zebrafish developmental atlas (Zebrahub) [Lange et al., 2023] to show how the incorporation of second-order LTRWs with memory, together with sequential information and RNA-velocity in StaVia, allows us to compute and visualize multi-lineage differentiation in atlases, and capture pathways that cannot be charted by other methods. We also demonstrate StaVia’s on a MERFISH dataset of the preoptic hypothalamus [Mofitt 2018], where StaVia integrates information of a cell’s spatial context with gene expression, to uncover sub-types and inter-cluster relationships that are missed when omitting the spatial information. Finally, a collection of developmental cell atlases are used to benchmark the Atlas View to other popular visualization methods, in which StaVia is the only method that successfully illustrates the temporal and lineage relationships in an 8 million cell dataset of mouse gastrulation [C. Qiu 2023]

## Results

### StaVia enables high-definition cartographic TI reconstruction across the entire single-cell atlas

StaVia is a graph-based TI framework designed to tackle challenges posed by atlas-scale data. It builds on our earlier Via method [Stassen 2021] which models cellular processes as a random walk with elements of laziness and teleportation across cluster graphs [Stassen 2020]. In StaVia we introduce higher-order walks with memory of cells’ previous states, integrated with cartographic views and enriched with information from available metadata (temporal or spatial), to reconstruct atlas-scale topologies coupled with automated predictions of diverse cell fates and their sequential specialization.

Advancing from Via and other TI methods, StaVia’s contributions are threefold. First, StaVia uses high-order LTRWs with memory to infer complex trajectories by relaying information about a cell’s previous states (**Fig. 1b**). This approach accurately pinpoints end-to-end differentiation paths and gene dynamics associated with a particular lineage. Forgoing the walk’s memory can obscure the distinction between the different pathways to cell fates in large atlases. Second, it allows flexible integration of data and metadata (e.g. time-series developmental labels from temporal atlases, spatial layout, gene/feature similarity and single-cell RNA-velocity) to compute pseudotimes, cell fates and lineage pathways (**Methods**) (**Fig. 1a**). Integrating available temporal data with the expression profiles allows us to stitch developmental points in a data driven manner. Spatial information is particularly challenging to incorporate when examining cellular landscapes based on their gene expression due to their highly non-linear nature. However, the microenvironments that cells occupy could provide valuable insight about their function. StaVia therefore provides a framework within which gene-expression and spatial information are jointly considered when charting the cellular landscape. Lastly, going beyond the common cluster graph visualization [Wolf 2019, Stassen 2021], StaVia generates an *Atlas View* that simultaneously illustrates complex chronological patterns and distinct phenotypic diversity, which has been challenging in current TI methods. Both the spatial arrangement of nodes and edges in StaVia’s high-resolution *Atlas View* (and its cluster graph), as well as their direction, connectivity, and weights, are guided synergistically by the results of the pseudotime, sequential metadata, and second-order LTRWs (**Methods**) **(Fig. 1a)**.

Using higher-order walks with memory is an unexplored feature in existing TI methods which typically rely on first-order random walks where prediction of future steps is independent of previous states (e.g., Palantir [Setty 2019], CellRank [Lange 2022], Via 1.0 [Stassen 2021]). When applied to reconstructing biological pathways, memoryless methods tend to encounter two types of problems which we explain by way of analogy to a faulty navigation system on a road trip from City A to B. The first issue occurs when memoryless methods suggest a cell passes through an unrelated intermediate population during development, akin to a GPS diverting us through an off-route City C. StaVia, uses higher-order LTRW with memory to act like an improved GPS that sense-checks directions, minimizing unnecessary detours, and ensuring a more accurate cell trajectory. Now imagine the road trip involves a critical turn at Point D but the GPS is so focused on the immediate road that it misses this turn. Analogously, some TI methods, due to an overemphasis on localized pathways, may fail to identify key transition states in a cell’s developmental journey. StaVia’s ‘memory’ feature is like an alert GPS that not only focuses on the road immediately ahead but also keeps track of the overall journey. It remembers where each cell has been and where it could be headed, making it less likely to miss critical turns (or transition states), and providing a more complete picture of the cell’s developmental journey. By integrating pseudotemporal forward biasing, RNA-velocity, and prior random walk state information (memory), StaVia’s higher-order LTRWs provide a more realistic prediction model of cell developmental pathways, enabling clear delineation of diverse lineages, transitional populations, and gene expression dynamics. **(Fig. 1b)**.

Our robustness analysis shows that adjusting the memory level has a predictable and gradual impact on lineage definitions, simplifying the optimization (See **Methods**, and **Fig. S5**). Generally, increasing emphasis on memory in the LTRWs yields pathways that emphasize the role of predecessors and remain inwardly focused. This translates to increased sensitivity to distinguishing related cell types and their gene expression dynamics **(Fig. 1b)**. Conversely, reducing memory helps explore poorly connected cell populations or those lacking precursors. The computational overhead from second-order LTRWs is minimal as they are conducted on the cluster graph level.

To generate StaVia’s cartographic *Atlas View*, we first create a single-cell embedding infused with second-order LTRW features learnt from the TI cluster graph (see **Methods**). Specifically, based on the presence of sequential data (e.g. data labeled with different time points), the single-cell graph can be sequentially augmented and refined accordingly. In the case of spatial data, spatial nearest neighbors are used to augment the gene-expression graph. Furthermore, prior to clustering and graph construction, the gene-expression is modified as the weighted average of a cell’s own cells and its spatial neighbors. We then use UMAP’s fuzzy simplicial set approach to align the high-dimensional LTRW feature space with the low-dimensional embedding (**Fig. 1a**) (See the definition of *LTRW feature space* in **Methods**). This single-cell embedding, a useful visualization in its own right, serves as the node layout for the Atlas View which arranges cell states and highlights edge connectivities (pathways) from the augmented single-cell graph using an edge bundling method based on kernel-density estimation (**Methods**). Directionality is projected on edges based on milestone pseudotime direction and RNA velocity. Note that the impact of high-order LTRWs with memory on the predicted end-to-end pathways can also readily be visualized in the Atlas View (**Fig. 1b**).

### StaVia captures a complete view of murine gastrulation

We employed StaVia on a scRNA-seq dataset of murine gastrulation [Pijuan-Sala et al., 2019], comprising 89,297 cells from stages E6.5 to E8.5 post-fertilization at quarter-day intervals. Previous trajectory analysis on this dataset required subsetting various lineages and analyzing them individually with manual curation in order to identify developmental trajectories of interest. In contrast, StaVia, by integrating higher-order LTRW with memory, RNA velocity, and time-series annotations (i.e., E6.5 to E8.5), accomplishes a holistic mapping of the entire atlas in a single run, accurately capturing relevant trajectories sans manual subsetting and curation (**Fig. 2-3**).

**Fig. 2.**
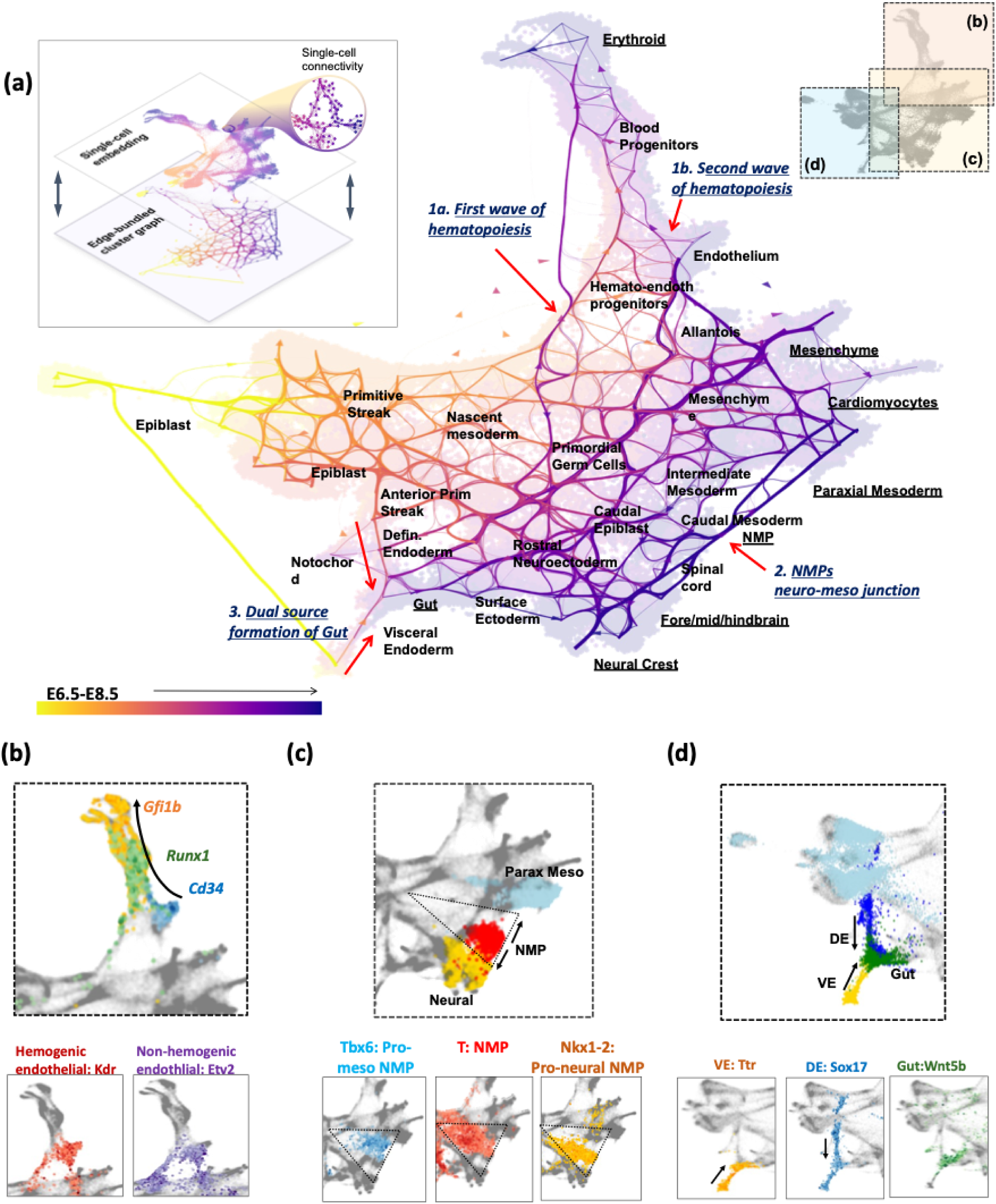
StaVia Atlas View of mouse gastrulation. (a) StaVia Atlas view of murine gastrulation, colored by known stage with edge directions inferred using a combination of RNA velocity and pseudotime. Root state automatically detected as epiblast E6.5. Autodetected terminal cell fates are underlined. (b) Sequential order of hemogenic endothelial cell differentiation. The black arrow is based on the edge direction of the hematopoietic branch in (a), and shows that Runx1 precedes the upregulation of GFi1b, which is a direct target of Runx1 and critically down-regulates endothelial markers to induce the EHT [Gao 2018, Thambyrajah 2016]. (c) NMP cells colored red reside between neural-yellow and paraxial mesoderm-blue (lhs) the zoomed in triangle of NMP cells express T brachyury. Of interest, the NMPs with a mesodermal tendency express Tbx6. NMPs with a more neural tendency express more Nkx1-2. (d) Zoom-in shows dual source of gut formation with Ttr positive cells at the Visceral Endoderm (VE), Sox17 expression for Definitive Endoderm (DE) and the gut itself expressing Wnt5b.

**Fig. 3.**
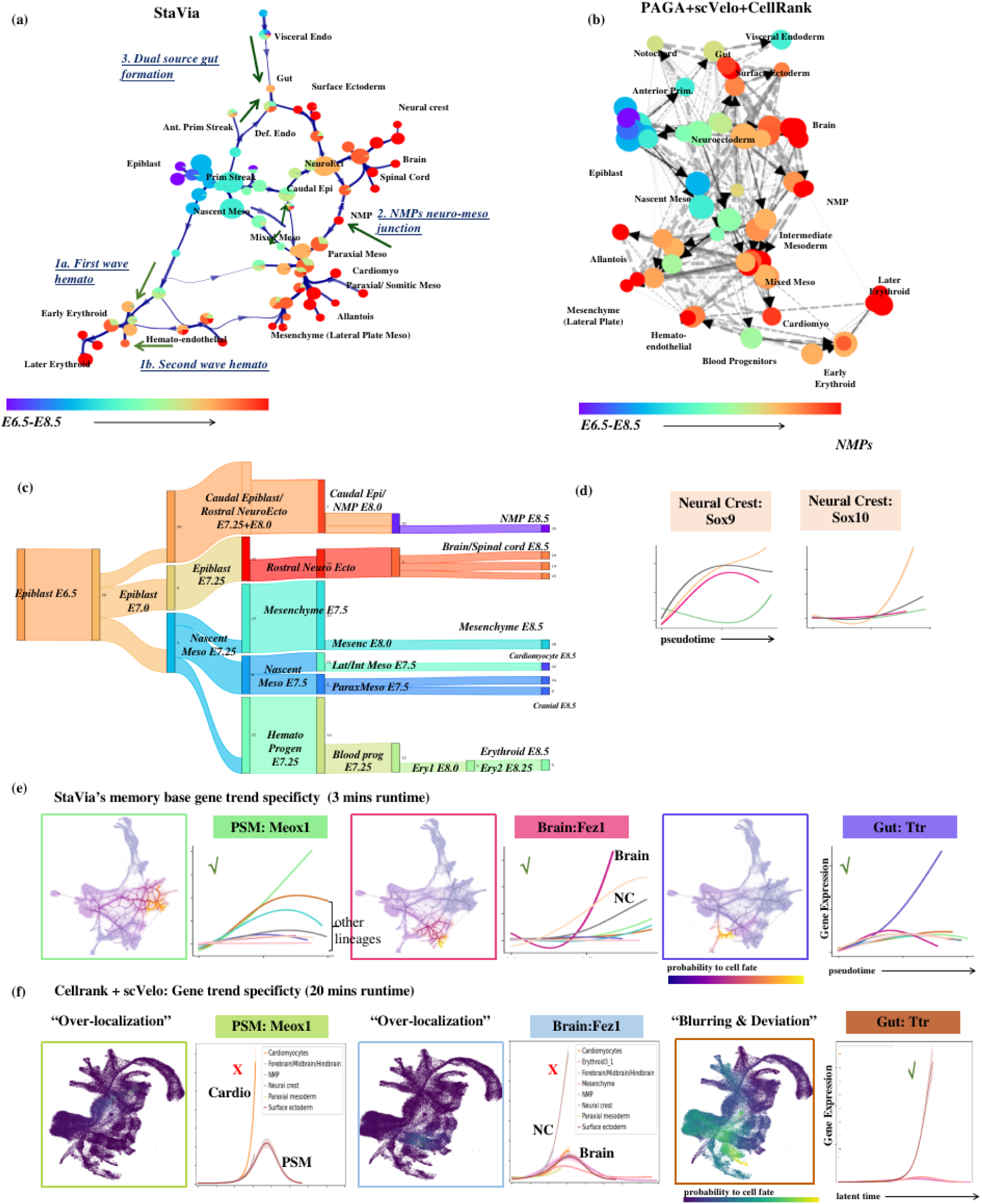
Comparison of TI graph structure and analysis. (a) StaVia cluster graph shows directed trajectory using a combination of scRNA-velocity and pseudotime. Colored by known stage Within the mesoderm, StaVia identifies cardiomyocytes, paraxial mesoderm, and mesenchymal cells; within the neuro-ectodermal branch: the surface ectoderm, brain and neural crest (NC); and arising from the visceral and definitive endoderm, the gut. (b) scVelo directed PAGA with a similar number of clusters and also using force-directed layout - lower visualized edges results in several disconnected clusters (c) Automatically predicted differentiation flow based on the cluster graph. (d) StaVia captures Sox9 upregulation preceding Sox10 in Neural Crest (NC) development. (e) StaVia end-to-end pathways from epiblast to cell fate for each germ layer. Each trend line corresponds to a lineage. The lineage of interest is highlighted by the color of the lineage-plot’s border and associated marker-gene. E.g. in StaVia the Brain lineage is dark pink. When the color of the marker gene matches the color of the upregulated trend line, it signals that the correct trend is inferred and merits a checkmark. (f) CellRank: The lineage-pathways to the Brain (light blue) exhibits is an example of where the pathway fails to detect transition states due to over-localization and the corresponding blue-lineage trend is not upregulated, warranting a cross-mark. The Gut pathway (brown trendline) is an example of deviation into unrelated intermediate states resulting in distinct pathways becoming blurred.

The Atlas View (**Fig. 2a**) and the cluster graph (**Fig. 3a**) visualizations created by StaVia illustrate the emergence of lineages within the entire dataset through a fanned-out structure that reflects the increasing separation between progressively specialized cells. The cluster graph (**Fig. 3a**), which forms the basis for the memory-infused second-order LTRW lineage probabilities as well as the layout and directionality of the Atlas View, captures the emergence of major lineages, their progressive separations towards cell fates (highlighted on the Atlas View **Fig. 2a** as underlined populations), and the correct placement of more subtle transition populations that exist at the boundaries of these major layers (e.g., neural mesodermal progenitors (NMPs)). Interestingly, both the cluster graph and Atlas View are uniquely able to show that the gut arises from the visceral and definitive endoderm. They also visually indicate two hematopoietic sources, the first being definitive erythroids from the primitive wave and the endothelial cells, which suggest the onset of the second wave **(Fig. 2a,b**, **Fig. 3a)**. This structure is not easily observed in other cluster graphs (e.g. PAGA **Fig. 3b**) or in higher resolution representations (e.g., UMAP, Phate), as evidenced by the comparative visualization analysis presented later (**Fig. 8**). See **Supplementary Note 2** for details of these three developmental patterns as shown by the StaVia cluster graph and Atlas View.

Next, we compared StaVia with a hybrid pipeline involving PAGA [Wolf 2019], scVelo [Bergen 2020] and CellRank [Lange 2022], a state-of-the-art method that combines gene-gene feature distances with directional information from RNA velocity for cell fate determination (See **Supplementary Note 1** for details on selection of methods for comparison). Comparing StaVia’s cluster graph with PAGA’s (using CellRank’s initial states and scVelo’s RNA velocity [Bergen 2020]) (**Fig. 3b) (**See **Supplementary Table S1** for detailed parameter setting), we observe that the PAGA-scVelo plot is visually difficult to interpret due to edge congestion that cannot easily be minimized. This is due to even conservative attempts of edge thresholding resulting in graph fragmentation. Importantly, the connectivity in the PAGA-scVelo plot misses key biological insights (e.g., lacks dual source of gut formation).

We subsequently compared the lineage probabilities from StaVia and CellRank towards different cell fates (**Fig. 3e-f**), with StaVia completing the TI computation in 3 minutes, compared to CellRank’s 20 minutes (**Supplementary Note 1**). Notably, the single-cell probabilistic lineages in CellRank do not capture the end-to-end pathways from epiblast, through transition states to final cell fates. In most cases for CellRank, the lineage probabilities (**Fig. 3b and Fig. S3** for all lineages) are either very localized to cells at the corresponding final cell fate with no indication of past states (NMP, brain, presomitic/paraxial mesoderm (PSM)), or are very diffuse detouring through the entire landscape (gut, cardiomyocyte) falsely suggesting that unrelated intermediate cells have a high likelihood of becoming these tissues. In contrast, the graph structure presented by StaVia (**Fig. 3a**) and its LTRW traversal using memory enables us to more unambiguously retrace how these lineages emerge (**Fig. 3a, 3c, 3e and Fig. S3**).

We note that the sequential pathways revealed by StaVia’s Atlas View are not recovered in CellRank which instead shows highly localized lineage likelihood without clear indication of originating or intermediate states (e.g. lineage probabilities localized to late paraxial mesoderm, brain in **Fig. 3f**, NMPs in **Fig. S3**) or very diffuse lineage likelihood (e.g. gut in **Fig. 3f** or cardiomyocyte lineage which spans the entire mesodermal trajectory in **Fig. S3**).

At higher memory levels, the brain lineage shows distinct elevation of *Fez1* and *Pax6*, crucial for neuroectoderm fate specification, neurogenesis, and forebrain patterning (**Fig. 3e, Fig. S4**) [Haedicke 2009, D. Duan 2013]. StaVia also reveals a noteworthy trend: *Sox9* expression precedes *Sox10* in neural crest (NC) precursors (**Fig. 3d**). This aligns with known data showing *Sox9*’s role in initiating premigratory NC cells, followed by *Sox10*, which fosters later NC development and cell emigration [Cheung & Briscoe 2003]. This *Sox9*-*Sox10* expression sequence is not captured by CellRank (**Fig. S3**).

### Introducing memory in random walks delineates end-to-end pathways in murine gastrulation

We next investigated how the incorporation of memory into random walks improves cell-fate mappings and their associated biological interpretation, by addressing the issue of too-localized or too-diffuse paths seen in current methods [Lange 2022]. In the murine gastrulation dataset (and later Zebrahub), we compared the lineage pathways obtained using a first-order and second-order LTRW with varying levels of memory (Mem=1 (no memory) to Mem=50). Lower memory values lead to more diffuse pathways on StaVia’s Atlas View **(Fig. 4 and Fig. S4)**, confounding analysis of temporal gene dynamics. In contrast, higher memory values successfully distinguish adjacent cell fates, as shown by the temporal gene expression of the NMP, paraxial mesoderm, neural, and neural crest cell fates at E8.5 (**Fig. 4a-b**).

**Fig. 4.**
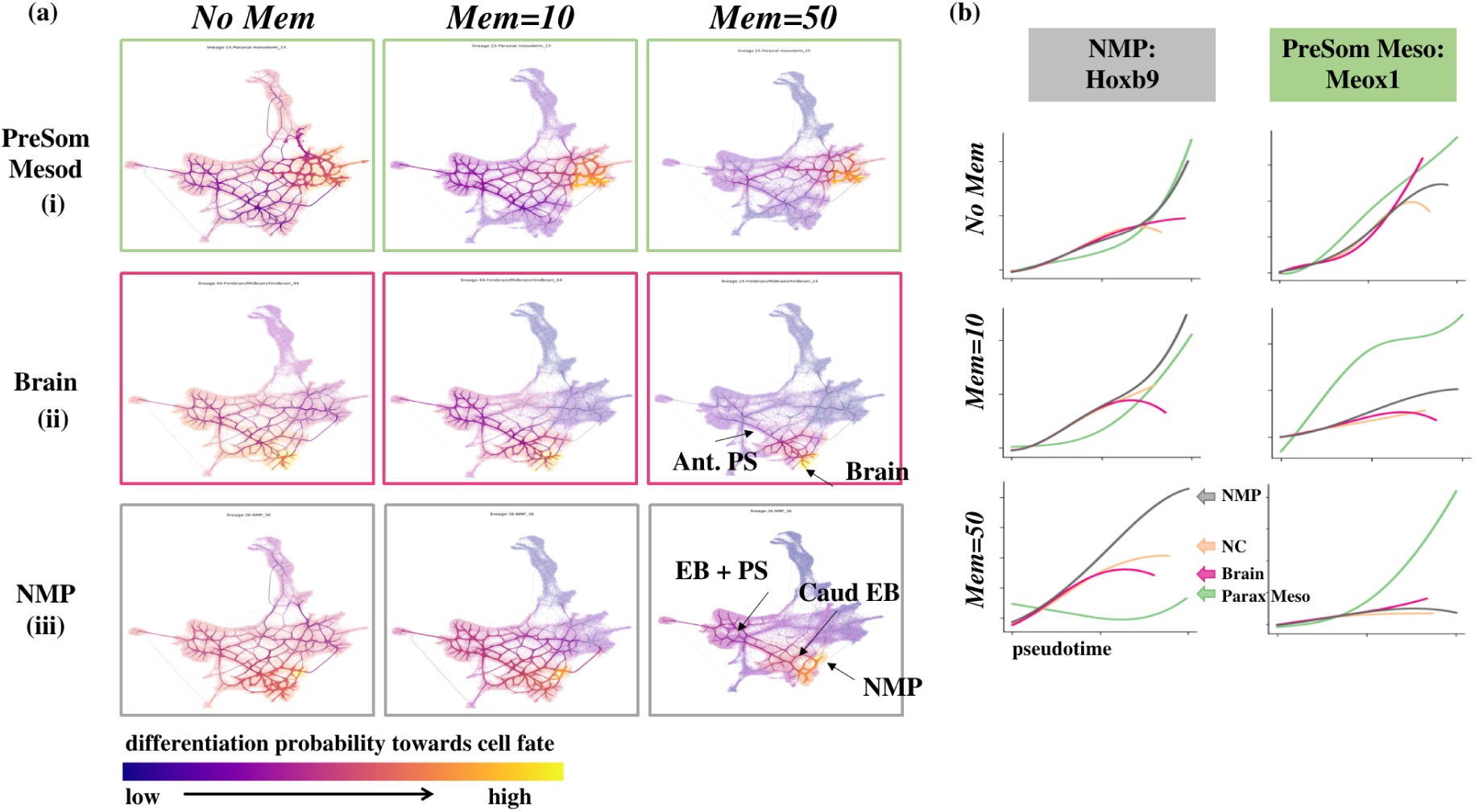
StaVia Memory impact on lineage paths. (a) Increasing memory mitigates too-diffuse pathways from epiblast towards specialized cell fates and consequently improves the associated gene trends specificity as shown in (b). Gene expression trends along pseudotime for lineages NMP (grey), PSM (green), Brain (pink) and Neural Crest (NC) (peach). Hoxb9 is an NMP marker and we expect the grey NMP gene expression to become comparatively more upregulated than the other three lineages. Similarly we expect Meox1 as a PSM marker to be comparatively more upregulated.

For the NMP lineage, nestled between the emerging PSM and neural cells, higher memory enables delineation of the sequential cell pathway from the epiblast to caudal epiblast, and then to the boundary of the mesodermal and neuronal lineages for the biopotent NMP cell fate (**Fig. 4a-iii**). In contrast, lower memory values tend to include unrelated cell populations. As a result, gene expression trends for NMP markers *Hoxb9* and *Nkx1.2* [Guibentif 2021] overlap for all these lineages at lower memory values, but at memory=50, the NMP lineage shows distinct expression elevation (**Fig. 4b, Fig. S4**). The benefits of the memory mechanism are also evident in the E8.5 brain cells **(Fig. 4a-ii)**, where higher memory more accurately shows the brain lineage deriving from the epiblast, followed by cells in the anterior pole of the primitive streak [Perea-Gomez 2015].

### StaVia displays holistic and high-resolution transcriptomic landscape of the full Zebrahub

We proceeded to leverage StaVia to probe the full Zebrahub, a recent comprehensive scRNA-sequencing time course atlas of 120,000 zebrafish embryonic cells [M. Lange 2023] (**Fig. 5a**). As current methods struggle to reconcile the extended temporal span and extensive cellular information of the entire 10-hpf to 10-dpf (hour/day post-fertilization) dataset, Lange et al., limited their study to the 10 to 24-hpf subset of cells (only 30% of the cells in the time-course study), omitting the peridermal and neuroectoderm lineages entirely. We show that StaVia successfully interrogates the complete dataset. Notably, Zebrahub’s neuroectoderm and periderm lineages are analyzed here for the first time with an example of the probabilistic pathway from each of these three layers (see the insets in **Fig. 5a)**.

**Fig. 5.**
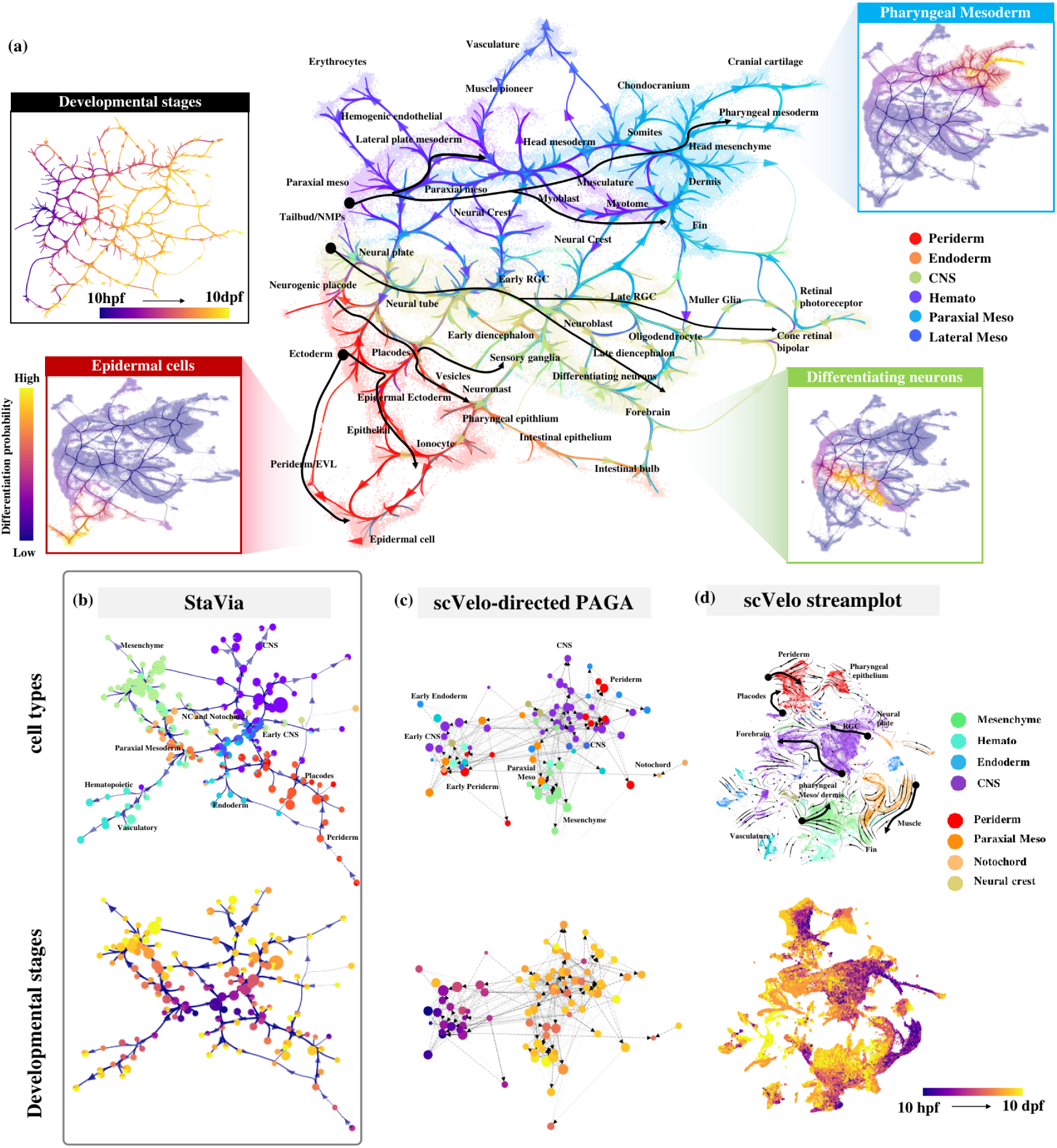
Zebrahub analysis by StaVia. (a) Atlas view of entire Zebrahub budstage to 10-dpf colored by germ layer. Black arrows highlight direction of differentiation indicated by Atlas edges for major lineages in the mesoderm, neuro-ectoderm and non-neuro ectoderm (Insets) StaVia end-to-end paths from bud to cell from from mesoderm, neuro and non-neural ectoderm show well delineated pathways as a result of higher order random walks. (b) StaVia directed cluster graph using scRNA-velocity and pseudotime colored by main tissue type (top) and known stage (bottom). Edge directions radiate outwards from the center. (c) scVelo-directed Paga constructed with a similar number of clusters as StaVia and also using a force-directed layout, shows a congested edge-layout with tissue-specific groups poorly separated and no clear direction. (d) scVelo streamplot on UMAP cannot mark the emergence of lineages as clearly as the edges in the Atlas view. The black arrows trace similar lineages to those highlighted in the Atlas, but do not transition through intermediate stages and often show conflicting direction

StaVia outperforms existing state-of-the-art methods, e.g., scVelo, in delineating the intricate cell differentiation trajectories. For instance, StaVia recapitulates that cross-talk between the major lineages (visualized as edges on both the Atlas View (**Fig. 5a**) and cluster graph (**Fig. 5b**)) is more prominent during earlier stages (e.g., neuro mesodermal progenitor pluripotent cell types like (NMPs) traverse two germ layers), and diminishes as cells become more specialized. Furthermore, the direction of differentiation is also more clearly captured by StaVia’s cluster graph (**Fig. 5b**) than CellRank-scVelo-directed PAGA representation (**Fig. 5c**) and scVelo’s streamplot (bold black arrows in **Fig. 5d)**. StaVia also detects more relevant late-stage cell fates (**Fig. 6b**, **7b**), as well as the gene trends that distinguish these lineages from each other (see Supplementary **Fig. S7-9** for lineage comparisons on all detected cell fates, where those missed by CellRank are manually provided to allow comparison), avoiding the pitfalls of missing transition states and inconsistent directionality occurring in the scVelo streamplot (**Fig. 5d**). Again, StaVia’s TI runtime is fast, taking 4 minutes, compared to 40 minutes in CellRank which misses several cell fates (**Fig. S6**).

**Fig. 6.**
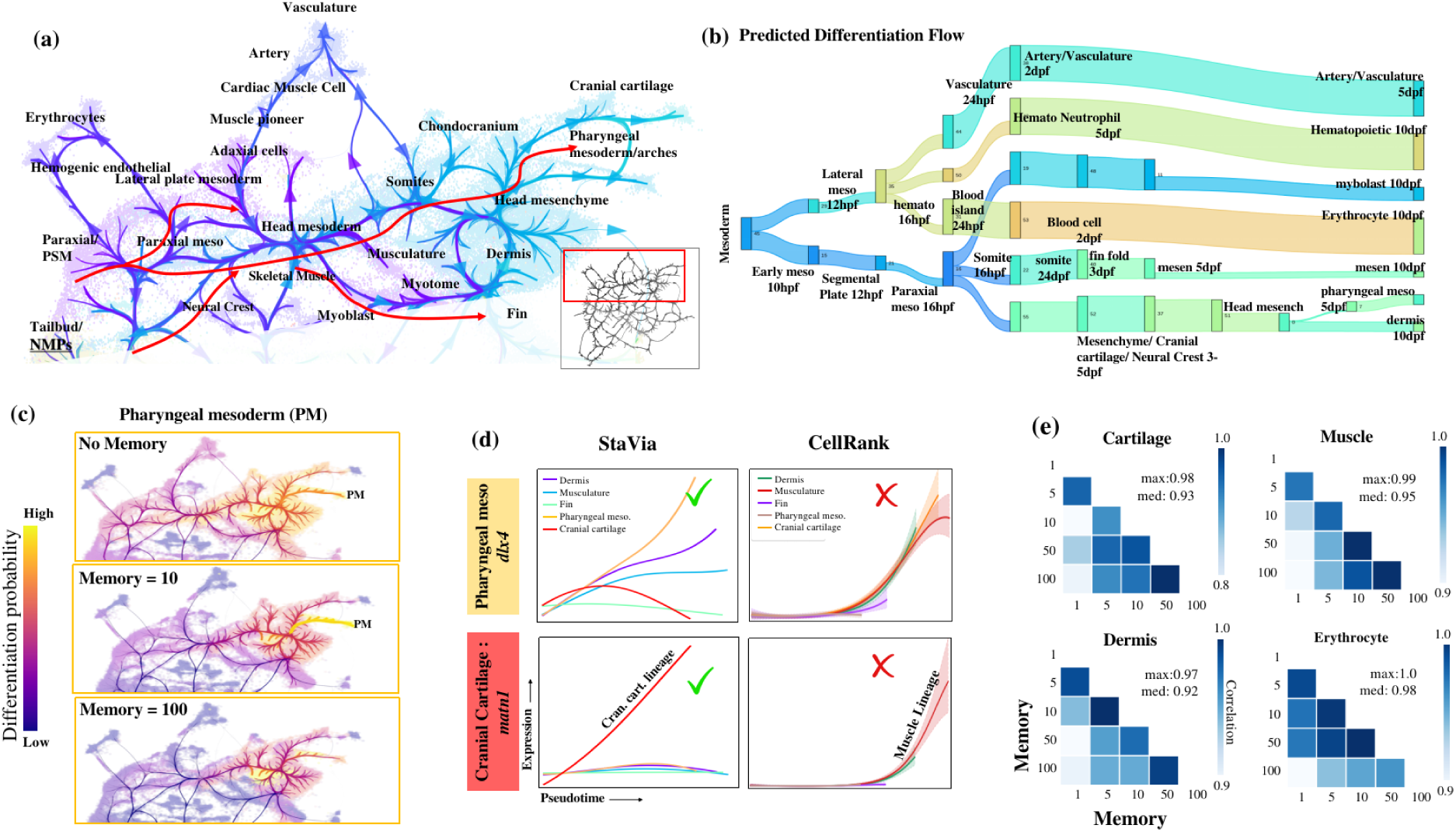
Mesoderm development revealed by StaVia. (a) Zoom-in of mesodermal lineage highlighting paths to pharyngeal mesoderm and musculature (b) automated predicted differentiation flow of detected mesodermal and hematopoietic cell fates from early mesoderm 10-hpf to 10-dpf (c) In StaVia, increasing memory shows clearer paths to cell fate of interest and avoids spillover into unrelated cell types. (d) Gene trends of mesoderm lineages for dlx4 (pharyngeal mesoderm marker) and matn1 (cranial cartilage marker) are correctly captured by StaVia, whereas CellRank’s lineages are incorrect (e.g. the muscle lineage upregulates matn1 in CellRank) (e) Correlation matrices for cartilage, muscle, dermis and erythrocyte lineage pathways at different values of memory 1 to 100 shows stability of analysis when changing memory (**Fig. S7** for all fates)

**Fig. 7.**
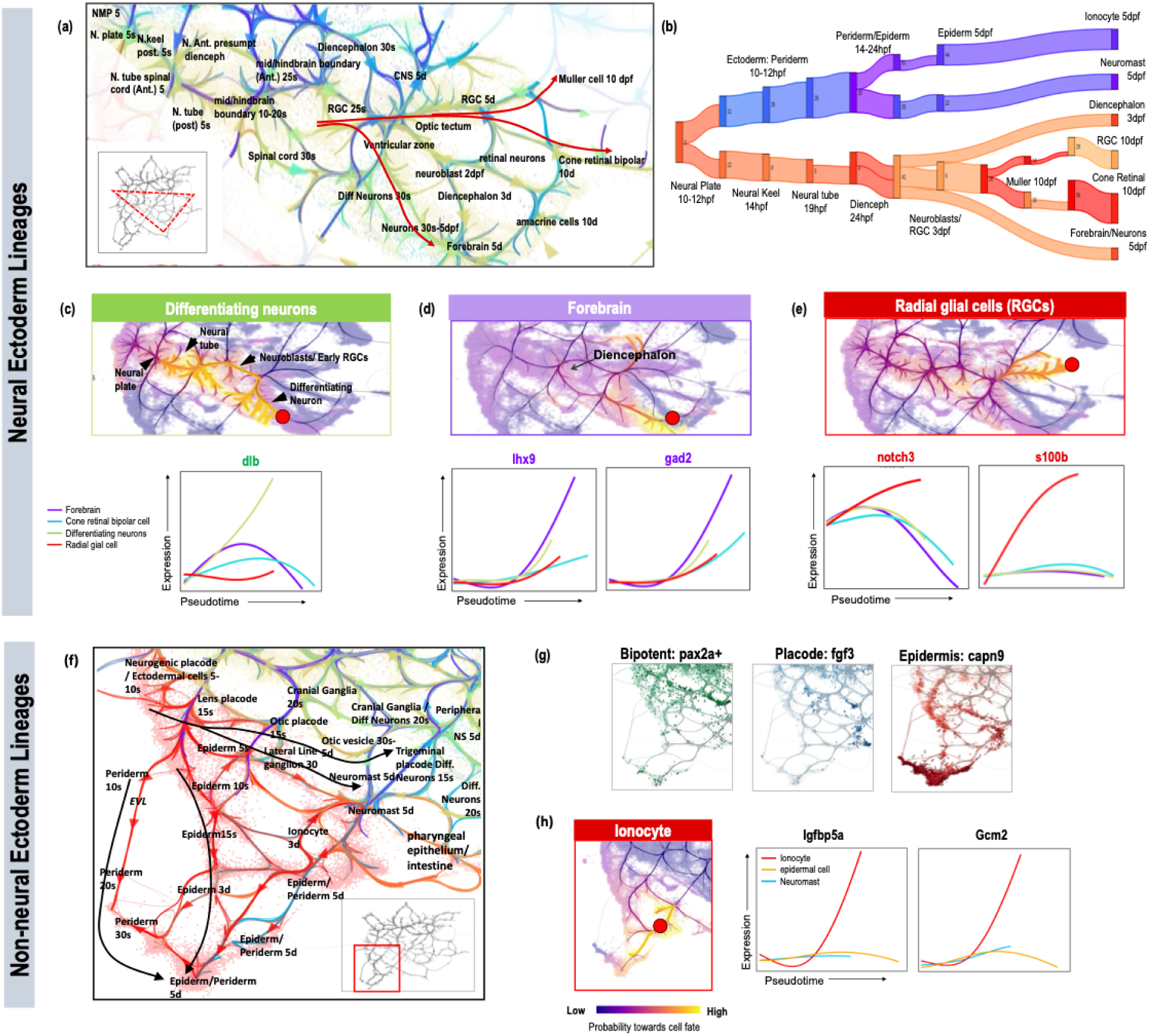
StaVia reveals complex ectodermal lineages. **(a)** Zoom-in Atlas View of neurulation from 5-hpf to 10-dpf. ‘s’ denotes somite-stage and ‘d’ is days post fertilization (dpf). (b) Predicted differentiation flow of non-neural ectoderm and neural fates. (c-d) End-to-end pathways from neural plate region at 10 hpf (5-somite) to (c) differentiating neurons and (d) forebrain (5-10 dpf). Accompanying gene-expression trends for neural lineages, shows upregulation of marker genes. (e) RGC end-to-end pathway and its marker gene expression trends. (f) Zoom-in Atlas View of non-neural ectoderm regions shows formation of bilayered epiderm and the differentiation of the placodes and their interactions with associated trigeminal neurons/ganglia. (g) pax2a expressed in the early epiderm and placode bipotent regions, fgf3 restricted to placodes, and capn9 concentrated on epidermal cells. (h) StaVia detects that the ionocyte fate (red dot) expresses more igfbp5a and gcm2 than other non-neuro ectoderm lineages.

### StaVia distinguishes multiple mesodermal pathways in Zebrahub

Again, StaVia uncovers a high-precision mesodermal differentiation flow (**Fig. 6a-b**) that cannot be recovered from the scVelo and PAGA maps. It accurately predicts that vascular and hematopoietic lineages are derived from the lateral plate mesoderm while revealing that the paraxial mesoderm gives rise to somitic cells, precursors to the dermis and cartilage. Critical to early embryonic development, bipotent NMPs, located at the early bifurcation of the mesoderm and neuroectoderm are identified **(Fig. 5a** and **Fig. 6a).** Again, this ability is attributable to the memory-centric graph traversal implemented in StaVia (See the impact of the memory on mesodermal differentiation analysis in **Fig. S10-11**).

Notably, StaVia accurately predicts that the pharyngeal arch is derived from the head mesoderm and the cranial neural crest [Mork & Crump 2015, Knight & Schilling 2006] (See the zoom-in Atlas View in **Fig. 6a**, differentiation flow in **Fig. 6b** and pathway in **Fig. 6c**)). The upregulation of *dlx* genes, as revealed by StaVia, marks the emergence of a pharyngeal population **(Fig. 6c)**. This stands in stark contrast to CellRank, which, lacking the memory mechanism, fails to distinguish *dlx* expression patterns across mesodermal lineages, resulting in a homogenized expression that masks true cell fate distinctions **(Fig. 6d, Supplementary Fig. S7** where cell fates missed by Cellrank are manually assigned to allow full comparison). Furthermore, StaVia identifies Matrilin *Matn* as a gene marker to distinguish the cranial cartilage from the pharyngeal arches **(Fig. 6eiii)**. This distinction is lost in CellRank (**Fig. 6d)**, which confounds the cartilage with smooth musculature. CellRank’s paths lack intermediate populations (**Fig. S7**), showing either fate-localized lineage probabilities or diffuse pathways (**Fig. S6**).

While increasing the memory for graph traversal can generally sharpen the specificity of lineage progression towards the desired cell fate (**Fig. 6c and Fig. S10-11** for other cell fates), excessive memory can constrain the pathway, as seen with the PM lineage at Mem=100 (**Fig. 6c**), underscoring the need for a balanced application of this parameter. Our stability analysis (**Fig. 6e and Fig. S5**) indicates that adjusting memory has a predictable and controllable impact. A heuristic correlating known time-series labels with pseudotime across memory values aids in determining the optimal memory range, ensuring the accuracy of the developmental inference by StaVia **(see Methods and Fig. S5**).

### StaVia elucidates neurulation sequence and differentiation of radial glia

We analyzed neuro-ectodermal lineages in Zebrahub (**Fig. 7a-e**) using StaVia, marking the first analysis of these cells which have otherwise been omitted in prior analyses. It identifies four distinct cell fates in the 5- and 10-dpf neuronal branches: forebrain cells, radial glia, and differentiating neuron, and cone retinal bipolar cells. This contrasts with CellRank’s identification of only the bipolar cells and radial glial cells (**Fig. S8** for the full set of cell fates). Crucially, StaVia successfully traces the neurulation sequence, from the neural plate through the neural tube cells’ progression to the diencephalon, and culminating in the mature forebrain neurons [Chatterjee & Li 2012] (**Fig. 7a-b**). Again, the use of memory enables us to identify gene expression trends specific to these cell fates. For instance, differentiating neurons are distinguished by upregulation of *Delta* genes (*dla/dlb*) specific to the subventricular zone **(Fig. 7c)**, whilst the mature post-mitotic neurons of the forebrain have elevated *lhx9* and *gad2* **(Fig. 7d)** CellRank’s diffuse probabilities of the neurons blur gene trends, confusing differentiating neurons with other ectodermal fates (**Fig. S8**).

The zoomed-in Atlas View (red arrows in **Fig. 7a**) and probabilistic pathways (**Fig. 7e**) uniquely highlight how the multipotent radial glial cells (RGCs) give rise to neurons and glia. They are characterized by activated *notch3* and *s100b* expression, indicative of early gliogenesis [H Li 2008, Dang 2006, Dimou 2014]. StaVia further captures the early RGCs are partially diverted to the differentiating neurons through a neuroblast sub-branch, while the other RGCs continue to the 10-dpf state where they differentiate (indicated by minor sub-branches) into Muller glia, oligodendrocytes or neurons (**Fig. 7a**).

### StaVia charts emergence of bilayered epidermis and placodes from Pax2+ field

StaVia’s Atlas View clearly separates the neural and non-neural ectoderm, enabling for the first time an unsupervised analysis of the Zebrahub non-neural lineages in the ectoderm (**Fig. 7e-g**). The identified edge connectivities capture how the ectodermal field of bipotent *Pax2+* cells give rise to both the *fgf3+* otic placode and *capn9+* epidermis [Ohyama 2006] (**Fig. 7f**).

The high-resolution edges of the Atlas view presents the emergence of otic placodes which later yield the otic vesicles **(Fig. 7e** black arrows). StaVia’s ability to capture localized details within a more global network is seen in the correct placement of the neuromasts (uniquely upregulating *fndc7a*) along the lateral line placode, with edges to the the neuronal cranial ganglia population known to innervate them. The formation of an early bilayered epidermis **(Fig. 7e)** from the extraembryonic enveloping layer (EVL)/periderm, and the inner basal epidermis is also detected by StaVia. The epidermal cells are identified by markers *capn9, anxa1b/c* in StaVia (**Fig. 7f, Fig. S9**), but in CellRank are indistinguishable from neuromasts and ionocytes due to diffuse lineage probabilities (**Fig. S9**).

Notably, StaVia detects two small cell fates each comprising less than 0.3% of the cell atlas. One comprises cells in the pharyngeal epithelial lining (located towards the lower right of **Fig. 7e**) which are formed by peridermal cells invading the pharyngeal cavity and subsequently expanding along the midline until the esophagus-gut boundary [Rosa 2019]. The second is the ionocyte cell fate (expressing *igfbp5a* and *gcm2* **Fig. 7h**), which are epithelial cells maintaining osmotic homeostasis. The ability to pinpoint these cell fates in the context of the entire dataset echoes the key strength of combining the Atlas View with the specificity of random walk memory.

### Systematic assessments of StaVia’s visualization

We systematically assessed the cartographic visualization performance of StaVia on six different single-cell transcriptomic datasets. One of these is the 8 Million cell mouse gastrula to pup atlas [C.Qiu 2023] **(Fig. 8e)** which was only computationally accessible to StaVia and UMAP - with StaVia being able to capture the developmental relationships in a more unified manner. We also benchmarked StaVia with commonly used single-cell visualization methods: UMAP, Phate, diffusion maps, principal component analysis (PCA), force-directed layout (ForceAtlas2 [Jacomy 2014]) and t-SNE [van der Maaten & Hinton 2008]. To facilitate comparison with other methods, we use the single-cell embedding generated by StaVia (prior to the edge integration step that creates the Atlas View), together with a set of five metrics that were adapted to account for the suitability of an embedding towards TI visualization. These metrics assess the ability to (1) convey progression and (2) separate lineages/distinguish cell types (**Fig. 8, Fig. S13a-b and Methods**).

**Fig. 8.**
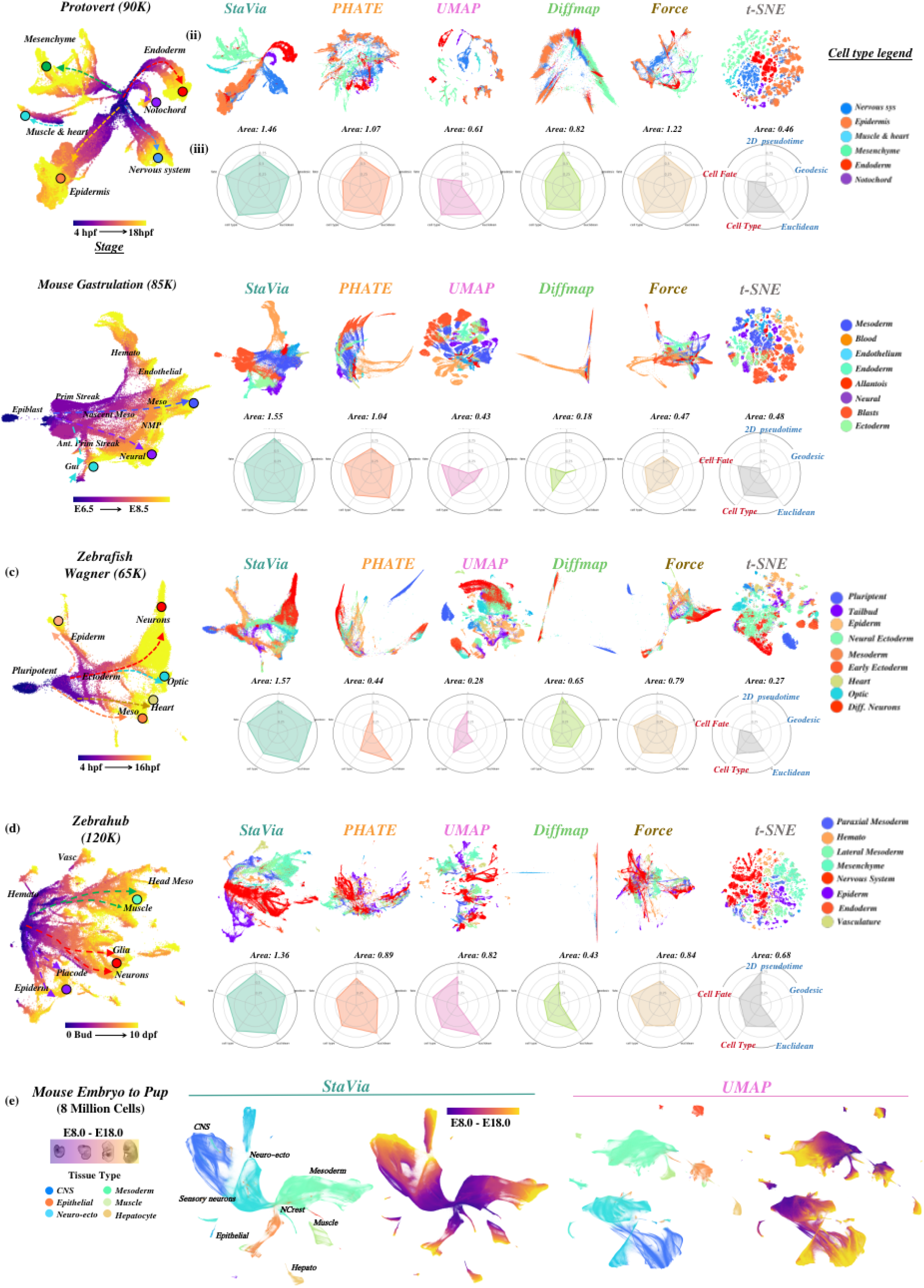
Comparison of Visualization methods. (a-d) (i) StaVia embedding colored by known experimental time (ii) Comparison of sc-embeddings generated by different methods colored by tissue type and (iii) radar plot scoring for each criteria. Read clockwise we have blue metrics (measuring sequential integrity) quantified by the correlation of the known time-series labels to the geodesic and euclidean distances from the root to other nodes in the embedded space, and also the 2D pseudotime uses the embedding as the input to StaVia rather than the original features/principal components (PCs) which would usually be used to compute the pseudotime for TI purposes. The second set of red metrics (separation of cell types/lineages) are measured by the cell-type F1-score when clustering the 2D input using the same number of clusters for all methods, and cell fate - measures how many fates are correctly detected by StaVia on the sc-embedding input rather than a higher dimensional input. (e) 8 Million cells of mouse gastrula to pup (E8.0 to E18.0) [C. Qiu 2023] by StaVia and UMAP colored by major tissue type and developmental stage. PHATE, t-SNE and force-directed layouts were attempted on this mega atlas but failed even after 24 hours of runtime and over 30 cores of parallel processing.

We demonstrated that StaVia’s single-cell embedding can consistently portray intuitive trajectory patterns, in accordance with the experimental time points (**Fig. 8a-d**). It also outperforms other competitive methods for faithful and robust TI visualization as evidenced by the five metrics (See the radar plot comparison in **Fig. 8a-d**). While UMAP and Phate are competitive methods for single-cell data visualization and have their respective strengths, we observed that Phate underperforms in being able to visually separate distinct cell types, although it significantly improves upon using selected diffusion components. Compared to Phate, UMAP scores well in delineating cell types, however suffers when it comes to visualizing connectivity between progenitor and progressively specialized cells. (See **Fig. S12 and Fig. S13** for all 5 datasets colored by stage and major tissue type across benchmarked visualizations) We also note that the superior TI visualization in StaVia does not compromise the computation speed, compared to other methods (**Fig. S12b** comparison of runtimes).

We also examined the impact of individual steps in StaVia on creating a TI compatible visualization (**Fig. S14** for detailed analysis of removing each step in the algorithm in turn and **Fig. S15-S16** for all 5 datasets colored by stage and tissue type). Removing sequential augmentation of the single-cell KNN graph and skipping the TI cluster-graph based initialization cause a significant drop in the ability to visualize progression. Keeping sequential augmentation but skipping the StaVia cluster graph initialization, which distills the underlying trajectory, also has a dramatic effect as quantified by the lower scores related to capturing the temporal progression as well as the visual outputs which appear more disjoint **(Fig. S14)**.

### Spatially aware cartography in StaVia captures relationships between cells based on both location and expression

Spatial omics have dramatically expanded our understanding of tissue architecture by mapping cells in their native environments, considering both their physical locations and gene expression profiles. Yet, it remains challenging to truly integrate this spatial information with gene expression profiles, resulting in analyses remaining purely in the transcriptomic domain with resulting cluster annotations and observations subsequently merely being visually projected back onto the spatial tissue locations. This makes integrative spatial and gene expression analysis non-trivial as it is now known that cell clusters or subtypes can exhibit stark contrasts in their distribution across a tissue slice, depending on their microenvironmental neighborhood, be it highly localized or dispersed.

While our examples have until now focused on temporally varying processes, we show that StaVia can also be used to investigate spatial datasets to understand cellular landscapes based on a combination of their expression levels as well as characteristics of their spatial “habitats”. As a proof of concept, we use the pre-optic mouse hypothalamus dataset based on MERFISH (multiplexed error-robust fluorescence in situ hybridization) [Moffitt 2018] to show that incorporating these spatial differences, in conjunction with gene expression, when clustering and capturing the connectivity-landscape elucidates differences in cell type and function. Here, StaVia’s graph construction leverages a recent concept [V. Singhal 2023] to recalibrate gene expression by considering a cell’s environment. Furthermore, StaVia also augments the gene-expression based KNN graph with spatial neighbors when establishing cluster connectivity (See Methods). Hence, the StaVia graph unifies gene expression with the spatial reality of the tissue.

In the hypothalamus dataset, this approach yields several key results: 1) the StaVia graph automatically generates and arranges clusters not purely based on their expression based cell type, but also their general tendency to occupy particular regions of the tissue section. While clusters themselves remain pure in terms of major cell classification, their cluster-level neighbors are often from a shared tissue ‘habitat’. The automated zoning of the tissue into neighborhoods of groups of clusters by the spatially aware cluster graph facilitates hypothesis generation and identification of potentially interesting sites in a tissue where different cell types are colocalized and potentially interact to yield location-specific functions. In the StaVia cluster graph **(Fig. 9a)** we use the nomenclature of tissue sub-regions [Paxinos & Franklin 2019] **(Fig. 9c)** to roughly guide the reader regarding the identified StaVia zones. Cells found in the lower section of the tissue slice are generally placed lower in the StaVia graph, whilst those towards the ACA and PVA are found in the top of the StaVia graph. 2) StaVia identifies sub-types that are missed when omitting spatial information. The excitatory neurons are separated into multiple subtypes that are located within their respective zones on the cluster graph but also express distinct DEGs **(Fig. 9b)**. For instance, the Oxytocin positive Excitatory cluster C42 is placed near the Ependymal cluster C9. *Oxt* neurons are known to be found near ependymal cells [Jurek & Neumann 2018], this linkage is not predicted when spatial information is left out of the computational analysis; both the oligodendrocyte and astrocyte population comprise two subtypes that present different spatial and DEG characteristics; and notably the sub-population originally annotated as “ambiguous” seems to actually comprise of both inhibitory neurons (Cluster C6 expressing *Gad1*) and Oligodendrocytes (C24 and C15 clusters, expressing *Mbp* and *Ermin*). We also use PAGA **(Fig. 9d)** as an example of a graph that does not relay very much spatial information and has a more limited set of sub-populations even when graph and clustering parameters are adjusted for, thus failing to achieve the results outlined above by StaVia.

**Fig. 9.**
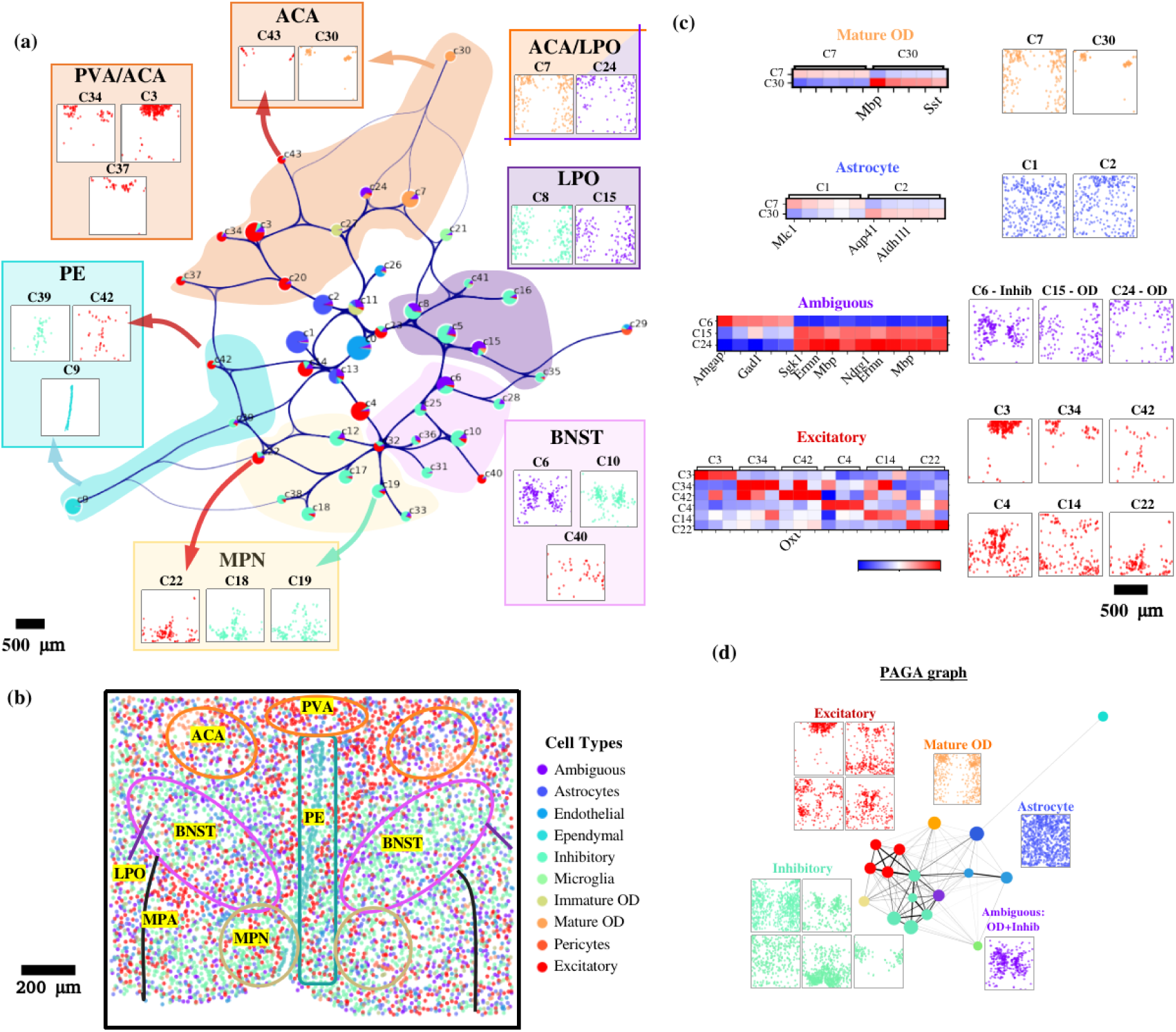
Spatially aware StaVia cartography for MERFISH data. (a) StaVia cluster graph of mouse hypothalamus preoptic region at bregma -0.289mm. Clusters located in similar zones share spatial ‘habitats’ Clusters are colored by cell type composition, all subplots share a legend for cell type coloration. Scatter plots placed near zones in the graph show the placement of cells in a cluster according to their spatial location on the tissue slice. (b) the top DEGs show clear differences in corresponding subpopulations (c) MERFISH tissue slice at -0.289mm colored by the cell type annotations originally found by Moffitt et al., (d) PAGA of the same cells without integrating spatial information. BNST, bed nucleus of the stria terminalis; MPN, medial preoptic nucleus PVA, paraventricular thalamic nucleus; ACA, anterior commissure; PE, periventricular hypothalamic nucleus LPO, lateral preoptic area MPA, medial preoptic area.

## Discussion

StaVia presents an advanced TI method integrated with a new visualization approach, tailored for cell atlases that encapsulate a high degree of complexity, be it diverse lineage representation, longitudinal temporal span, non-linear spatial layout, or sheer sample size. A salient feature of StaVia is the implementation of the second-order random walks with memory, vital to delineating the intricate end-to-end pathways of multiple lineages in the entire differentiation process.

We demonstrated that without higher-order random walks with memory (by comparison to CellRank and by removing memory), lineage pathways run into two main issues: deviating into unrelated intermediate populations which entangles pathways and hinders lineage specific insights, or becoming so myopic in search paths that transition states are overlooked. The usefulness of StaVia’s random walks with memory to overcome these challenges without resorting to manual subsetting of stages/populations that would otherwise rob the atlas of its unique scale and perspective, was demonstrated on the mouse gastrulation dataset. StaVia identified sequential transitions in hematopoiesis with new insights into hemogenic endothelial differentiation, NMP bipotency and dual-source gut formation that are not detected by other methods without manual curation and subsetting of the atlas.

StaVia also facilitated the collective analysis of all cells in the recent Zebrahub atlas for the first time. The use of memory together with automated integration of time-series information revealed insights into the bilayered epiderm formation, placode development and the differentiation of glial cells during neurulation. Moreover, StaVia’s runtime for retrieving lineage pathways is competitive, requiring a few minutes in comparison to CellRank which needed more than 30 minutes on the same dataset using the same hardware. Efficient processing and runtimes aid in the discovery process by allowing the analysis to be probed across parameters on a collective atlas level without requiring access to immense computational resources.

Current state-of-the-art methods (e.g. the scVelo-directed PAGA graphs, and the RNA-velocity stream plots) quickly become congested in terms of edges or streamlines and struggle to convey biological transitions. For effective visualization of complex trajectories at an atlas scale, it is imperative to establish a linkage between the overall network structure and the fine-grained transcriptomic signature [M.M.Li 2022]. StaVia’s Atlas View does this with its high edge resolution and TI-based spatial layout of cells, providing perspective on biological chronology while preserving spatial proximity of similar cells, being uniquely able to visualize development in the 8 Million cell gastrulation atlas.

As shown on the preoptic MERFISH dataset, StaVia’s framework can integrate spatial information of cells on tissues, offering perhaps the first cartographic approach to spatial transcriptomics data that conveys both location and gene-expression based similarities of cell types. This revealed several sub populations that had originally eluded separation based purely on gene-expression and also showed intra-cluster relationships between cell types based on physical location. As we anticipate an increase in the creation of cell atlases with both temporal and spatial emphases, StaVia’s capabilities in delineating and visualizing cellular trajectories in large-scale and complex datasets could spearhead more large-scale bioinformatic strategies that enable a more comprehensive understanding of cellular differentiation, lineage trajectories, and disease progression.

## Methods

### Key steps in the TI Algorithm and Visualization in StaVia

StaVia is built upon our earlier work of Via [Stassen et al., 2021] that models the cellular process as a modified random walk, called LTRW, transversing the cluster graph computed by a data-driven community-detection algorithm [Stassen et al., 2020]. This model incorporates elements of ‘laziness’ (staying at the current state) and ‘teleportation’ (jumping to any other state), with predefined probabilities. Pseudotime and graph directionality are then calculated based on state hitting times, and refined with Markov chain Monte Carlo (MCMC) simulations. Here below are the key elements and steps relevant to the StaVia framework:

1. *<u>Represent single-cell data by a sequentially augmented graph:</u>* The first step is to represent the single-cell data by a single-cell-KNN (scKNN) graph using the hierarchical navigable small world algorithm [Stassen et al., 2020]. Subsequently, if sequential (temporal) data (e.g. data at different time points) is provided, then additional edges between cells in adjacent sequential groups are added. Edges between cells that are more than *t_threshold* can also be optionally removed. In the case of spatial data the input gene expression is first modified to a weighted average of a cell’s own cells and that of its spatial neighbors. The construction of a scKNN is done in this new gene expression space (following PCA) and then augmented by spatially adjacent neighbors added to the scKNN based on spatial proximity.
2. *<u>Build the cluster graph:</u>* Following this, a cluster graph is constructed where nodes are PARC-based clusters of single cells. These groups of nodes can also be pre-defined by the user. Similar to Via 1.0 a pseudotime based on LTRW is first computed and the edges are accordingly forward biased. In StaVia, when available, the edge directions are also determined by the scRNA velocity. The edge-weighting and direction given by scRNA velocity versus pseudotime is controlled by a user-defined parameter (set as 0.5, i.e. 50/50 weight, by default). Start states are predicted based on the absorption probability (when scRNA velocity is available) or defined by the user. Terminal states are computed similarly to Via 1.0 using node degree and connectivity properties.
3. *<u>Compute the lineage pathways:</u>* Pathways (from the root to the terminals) are computed using second-order LTRWs with memory conducted on a forward-biased (directed and weighted) cluster graph. These give us a collection of simulated second-order random walks that describe the probabilistic pathways. The *memory_parameter* controls the weighting-multiplier used on node edges.
4. *<u>Construct an edgebundled cluster graph:</u>* The cluster graph’s node layout is computed using the fruchterman-reingold method. Despite the relatively modest number of clusters, edge congestion quickly arises with pruning of edges being a suboptimal way of reducing clutter. We therefore visualize edges using an edge bundling technique based on kernel density estimation (KDE) which transforms the graph into a density map and then moves edges towards the local density maxima to form bundles [Hurter 2012 and CuBU 2016].
5. *<u>Construct a single-cell embedding for Atlas View:</u>* The underlying single-cell embedding relies on UMAP’s implementation of minimizing the fuzz-set cross-entropy between the high and low dimensional representation. In our case, the high dimensional representations (typically 50-100 features) are the Node2Vec [Grover & Leskovec 2016] features learned from the second-order LTRWs on the sequentially augmented scKNN graph which tend to preserve global relationships better than principal components (**Supplementary Fig. S14** analysis of individual steps in StaVia visualization) The cell-cell neighborhood used in the cost function computation is based on the sequentially augmented scKNN graph. The initialization of the single-cell embedding is based on the layout of the forward-biased TI cluster graph.
6. *<u>Generate a complete Atlas View:</u>* The single-cell 2-D embedding layout, as described above, is clustered using kmeans (set to 150-1000 clusters depending on the desired granularity). A cluster graph is formed using the augmented sc-KNN graph and the edges are bundled using KDE. These are overlaid on the TI-cluster graph initialized single-cell embedding from Step 4 to create the complete Atlas View. The bundled edges greatly aid in visually sharpening the spatial density of edges and emphasizing high traffic edge patterns.

### Second-order LTRW in StaVia

The concept of second-order random walks have been used previously to define search neighborhoods in feature representations of networks which can subsequently be used for classification tasks [Grover & Leskovec 2016, Liu & Krishnan 2021, Liu 2023]. We extend this idea to TI computation in StaVia. Here, we use a fast implementation of the node2vec algorithm [R.Liu et al., 2023] to compute second-order walks on the directed cluster graph (used for lineage probability predictions and pseudotime) and to compute a high-dimensional feature representation of the sequentially augmented network. These node2vec graph-representation components are used instead of principal components when optimizing UMAP’s fuzzy simplicial set’s cost function.

The cluster graph constructed in StaVia is defined as a weighted connected graph G (V, E, W) with a vertex set V of n vertices (or nodes), i.e. 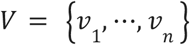 and an edge set E, i.e. a set of ordered pairs of distinct nodes. W is an *n* ×*n* weight matrix that describes a set of edge weights between node i and j, *w_ij_* ≥0 are assigned to the edges (*v_i_*, *v_j_*).

Assume the walker is currently on node *v_curr_* and has neighborhood *N_v_* comprising three neighbors (*v_m_*, *v_n_*, *v_o_*) **(Supplementary Fig. 18a)**. In the first order case the transition probability is given by

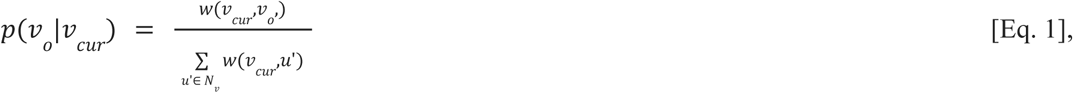

where the probability is only conditioned on the current state. However, in the second order random walk framework we adapt for StaVia’s lineage probability computations **(Supplementary Fig. 18b)**, a bias factor *alpha α* **≤1** is applied to reweight edges depending on the previous state such that neighbors of the current node that are not neighbors of the previous node are considered “out-edges”. A node that is a mutual neighbor of the current and previous nodes is an inward edge, with the return-edge being the case when the next node returns to the previous one. When *α* = 1 this system reverts to the first order case. The original node2vec algorithm applies an additional biasing parameter for the return-edge to discourage getting stuck in a loop that returns to the previous state. However, in our case, since we have a forward-biased weighted graph that already suppresses reverse behaviour against the pseudotime, this additional biasing is not required. The transition probability in the second order case is now given by

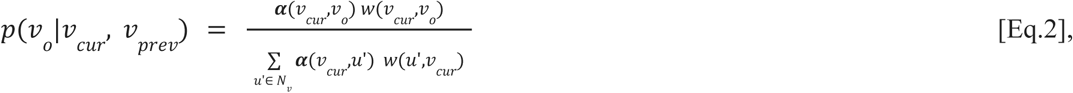

which generalizes to

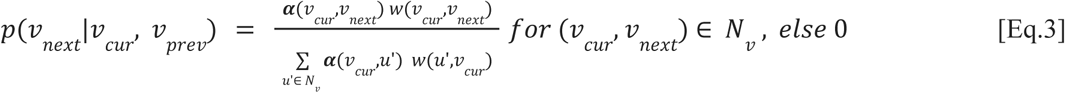

and it has bias factor *α*, given by

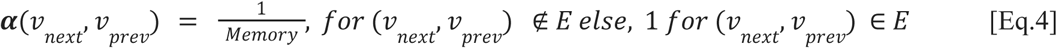

Stability of the TI when changing the memory parameter and a short guide to selecting a suitable range is described below (See the **Robustness analysis of memory parameter** and in **Fig. S5**).

### Node feature representation

The biasing framework described in Eq.3-4, adapted from Node2Vec, was originally used to sample node neighborhoods used to compute the cost function for node feature representations. Random walk based approaches to node embeddings aim to maximize the probability of reconstructing the neighborhood *N_s_* (*u*) of any node *u* in a graph based on some sampling strategy *S* conditioned on the feature representation *f* [Grover & Lescovec 2016]

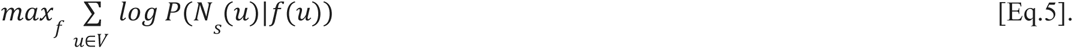

When computing the Atlas embedding, we use these graph representation features instead of principal components (PCs) to calculate the fuzzy-set cross entropy between the sc-KNN sequentially augmented graph and the 1-skeleton graph obtained from the embedding method.

### Kernel density estimation (KDE)-based edge bundling

The graph bundling for the cluster graphs and Atlas view uses a kernel density estimation based method [Hurter 2012, Cottam 2014 and CuBU 2016]. Combining this with StaVia’s single-cell atlas embeddings aids in visually summarizing the edge density and highlighting pathways based on their traffic (cell-cell interactions). Briefly, the KDE edge bundling is an iterative algorithm that repeats the following set of steps on a graph drawing: first convolve the edges with a kernel to construct a density map. The density is a measure of the number of edges at that particular location in space. Next, compute the gradient of the density map *Δρ* and advect points 𝐱∈𝐺 in the direction of *Δρ* and do Laplacian filtering to smooth the edges. Repeat these steps, reducing the kernel bandwidth on each iteration. The effect will be to sharpen the density such that straight-lined unbundled edges will be drawn as tightly bundled curves.

Given a graph drawing 𝐺⊂𝑅^2^ with edges 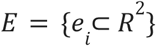 where 𝐱∈𝐺, the density map is given by

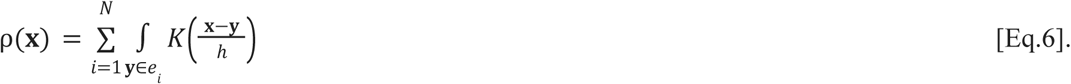

The kernel used here is the Epanechnikov kernel

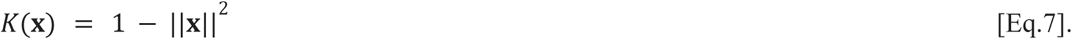

The bandwidth is reduced by a factor λ at each iteration, such that on the *n^th^* step it will be 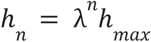

### Robustness analysis of memory parameter

We have investigated the utility of incorporating random walks with memory in the context of both mouse and zebrafish gastrulation, whereby gradually increasing the level of memory in the random walk improves end-to-end pathway mapping and the specificity of gene trends associated with the emergence of lineages. However, at very high levels of memory (e.g. memory =100), some pathways can be too restrictive. We show that it is possible to narrow down or gauge an optimal range for the memory parameter by correlating the known experimental times to the inferred pseudotime (computed at different memory values). For Zebrahub and Mouse gastrulation, the correlation increases with memory, remaining elevated for an interval before decreasing at higher levels of memory **Supplementary Fig. S5b-c**. To show that the change incurred by memory is gradual and behaves in a stable manner, we also present a correlation analysis in **Supplementary Fig. S5a** of lineage probabilities for cell fates in the Zebrhub dataset at memory values {1,5,10,50,100}, where a value of 1 signifies no memory. These show that for a wide range of values, the analysis is highly correlated.

### Metrics for quantitative analysis of visualizations

The metrics fall into either structural or cell type measures. Structural metrics to assess how well a method visualization sequential information and progression include standard pearson correlation 𝑟(𝑥, 𝑦) between :

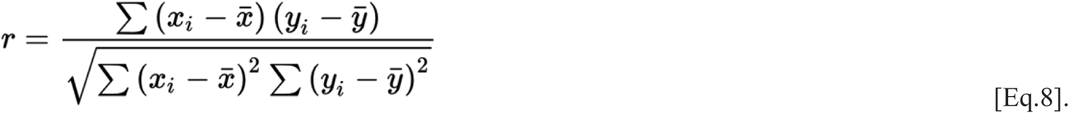

- *<u>2D-pseudotime:</u>* Time series label and StaVia-pseudotime, where StaVia-pseudotime is the inferred pseudotime by StaVia when the 2D embedding is given as the input to StaVia.
- *<u>Geodesic:</u>* Time series label and geodesic distance on the embedding from root to cells. The geodesic distance *<u>d</u>^geo^*(*u*, *v*) between two nodes 𝑢, 𝑣 on a weighted graph is the minimum sum of weights across all the paths connecting *u* and *v*.
- *<u>Euclidean:</u>* Time series labels and euclidean distance on the embedding from root to cells. The euclidean distance *<u>d</u>^Euc^*(*<u>u</u>*, *v*) between two cells 𝑢, 𝑣 whose coordinates are given by the 2Dembedding,

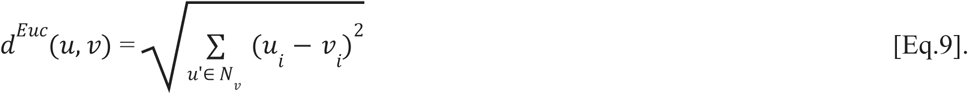

On the other hands, the metrics that assess how well a visualization method captures lineage divergence and respects cell type separation include:

- *<u>Cell Fate:</u>* Cell fate detection when running StaVia on the embedding. StaVia’s automatic cell fate detection is applied to all embeddings (the same root state is provided in all cases). We compute the F1-score of detected cell fates with reference to expected cell fates (corresponding to the later stages of the dataset)
- *<u>Cell Type:</u>* Use k-means clustering on the 2D embedding at a fixed number of clusters for all embeddings. For each dataset, the number of clusters is set to 5 clusters more than the number of given coarse-level annotations resulting in typically around 15-20 clusters. Calculate the

F1-score using the scoring method applied in Stassen 2020 which assigns each cluster a majority reference population, aggregates the clusters assigned to said reference population and calculates the one-vs-all F1-score for each reference population. The mean score across reference populations is reported which avoids the issue of larger cell populations dominating the score. This approach prevents punishing a method for splitting a cell type into multiple clusters, which may well be the case since the annotations are coarse and would not necessarily capture subtypes. Since all methods are given the same number of k-clusters, it is still a fair comparison.

### Spatially aware cartography construction in StaVia

Our strategy has two key elements which play a role in embedding spatial information into the cartography. The first element uses a concept [V. Singhal 2023] that recomputes the gene expression as a weighted average of a cell’s own expression plus that of its spatial neighbors. The second component of the spatial integration is to augment the single-cell gene-expression-based KNN-graph with neighbors found in the spatial domain before computing inter-cluster connectivity. The average spatial location of clusters is subsequently used to initialize the layout of the StaVia graph.

### Pre-processing datasets

#### Mouse Gastrulation (Pijuan Sala 2019)

Scvelo’s filter_and_normalize function is used on the raw spliced and unspliced genes. Only genes (both in spliced and unspliced counts) that are expressed in 20 or more cells are retained, resulting in a matrix with 10,766 genes across 89,267 cells. Each cell is normalized by the counts over all its genes. The last step is to log normalize the counts before PCA. The velocities are computed using scvelo’s stochastic mode as the dynamic mode is prohibitively slow for large datasets. PCA done on the full filtered gene set.

#### Zebrahub (Lange 2023)

single-cell sequencing of 120,437 cells from zebrafish embryonic development across 10 timepoints from 10-hpf to 10-dpf. Cells expressing fewer than 200 genes and genes expressed by fewer than 5 cells were removed. Each cell is normalized by the counts over all its genes, followed by log normalization. The top 5000 highly variable genes across 120,437 are used for PCA. Due to the computational demands of Scvelo for large datasets, we use the velocity matrix made publicly available by Zebrahub which was computed using scVelo. Cell annotations were combined from the datasets for individual timepoints.

#### Ascidian Protovert (Cao 2019)

Early phases of embryogenesis of ascidian protovertebrate with sequentially staged Ciona embryos, from gastrulation at the 110-cell stage to neurula and larval stage. Individual h5 matrices for each timepoint are concatenated into an Anndata object containing 90,579 cells. Standard gene filtering (min_cells=5, min_counts=10) is done using scanpy and each cell is normalized by the counts over its genes followed by log normalization. Top 2000 highly variable genes are used towards PCA. Root cell is user defined based on time-stamps as stage-1 epidermis cell.

#### Zebrafish (Wagner 2018)

63273 cells across 7 timepoints from the first 24 hours of zebrafish embryo Same preprocessing as Ascidian Protovert data. Top 2000 highly variable genes are used towards PCA.

#### Mouse Neuron (La Manno 2021)

292495 cells from embryonic mouse brain from stage E8-E18. Same preprocessing as Ascidian protovert. Scvelo is used to compute the scRNA velocity. Top 2000 highly variable genes are used towards PCA.

##### MERFISH (Moffitt 2018)

MERFISH of 12 slices of mouse preoptic region 1.8mm x 1.8mm x 0.1um thick of 160 genes chosen based on scRNA-seq marker gene panel and known functional gene panel. Fig. 9 is 6500 cells from a naive female mouse slice at -0.289mm bregma. Analysis of slices from other bregma shows similar spatially aware StaVia graphs and clustering tendencies.

##### Mouse Pup (Qiu 2023)

Three-level single cell transcriptional profiling by combinatorial indexing (sci-RNA-seq3) profiling over 8 million nuclei from 83 staged embryos spanning late gastrulation (E8.0) to the end of gastrulation at E18.75, with 2-hr temporal resolution during somitogenesis and 6-hr resolution through to birth.

<u>TI Parameters</u> for StaVia, Paga, CellRank and <u>Visualization parameters</u> for all methods are provided in Supplementary Table S1 and Table S2 with number of KNN and PCs consistent for each method and key parameters highlighted where changed from default in order to improve results. No batch-correction based on experimental times was performed for the datasets during TI analysis or visualization benchmarking. The UMAP embeddings used to present Cellrank’s TI results use the UMAPs resulting from batch-corrected PCs used in the Zebrahub and Sala publications.

## Data Availability

Raw and processed data for Zebrahub is available at [https://zebrahub.ds.czbiohub.org/]. Mouse gastrulation data is available at https://bioconductor.org/packages/release/data/experiment/html/MouseGastrulationData.html. Data for Ascidian Protovert is available at https://www.ncbi.nlm.nih.gov/geo/query/acc.cgi?acc=GSE131155 Data for Zebrafish Wagner is available at https://kleintools.hms.harvard.edu/paper_websites/wagner_zebrafish_timecourse2018/mainpage.html. Data for Mouse Neuron data is available at http://mousebrain.org/development/downloads.html. Processed Anndata h5ad files used in this paper for the datasets above are also available at https://github.com/ShobiStassen/VIA

## Code Availability

StaVia is available as a pip installable python library “pyVIA” with tutorials and sample data available on https://github.com/ShobiStassen/VIA, https://pyvia.readthedocs.io/en/latest/index.html. Gallery of Atlas Views for more datasets (including the 8 million cell gastrula to mouse pup from Qiu 2023) are available at https://pyvia.readthedocs.io/en/latest/Mouse_to_pup_atlas.html.

## Supplementary Materials

See accompanying attachment for the following figures, notes and tables

## List of Figures

Supplementary Fig. S1. Comparing TI graph structure of Via2.0 and CellRank+PAGA

Supplementary Fig. S2. Lineage paths and gene trends for cell fates in StaVia using memory

Supplementary Fig. S3. Lineage paths and gene trends comparison between StaVia and CellRank

Supplementary Fig. S4. StaVia Memory helps distinguish gene trends near the NMP lineages

Supplementary Fig. S5. Stability analysis for memory

Supplementary Fig. S6. CellRank lineage probabilities not improved by projecting on Atlas View

Supplementary Fig. S7. Zebrahub mesoderm: Comparison of lineage paths and gene specificity trends

Supplementary Fig. S8. Zebrahub neural ectoderm: Comparison of lineage paths and gene trends

Supplementary Fig. S9. Zebrahub non-neural ectoderm: Comparison of lineage paths and gene trends.

Supplementary Fig S10. Impact of Memory on Mesodermal Lineage Probabilities

Supplementary Fig S11. Impact of Memory on Mesodermal Gene Trends (Zebrahub)

Supplementary Fig. S12 Comparison of visualization methods on different time-series RNA-seq datasets (cells colored by tissue type)

Supplementary Fig. S13 Comparison of visualization methods on different time-series RNA-seq datasets (cells colored by developmental stage)

Supplementary Fig. S14 Impact of key steps in StaVia Atlas View embedding

Supplementary Fig. S15 Impact of Steps in StaVia Embedding (cells colored by developmental stage)

Supplementary Fig. S16 Impact of Steps in StaVia Embedding (cells colored by tissue type)

Supplementary Fig. S17 Impact of Steps in StaVia Embedding (Radar plots)

## List of Tables and Notes

Supplementary Table 1 for parameters used in TI

Supplementary Table 2 for parameters used for single-cell embeddings

Supplementary Note 1: Selection of TI methods used for comparison

## Acknowledgements

This work was funded by the Research Grants Council of the Hong Kong Special Administrative Region of China (grant nos. 17208918, RFS2021-7S06, and C7047-16G).

## Author Information

### Contributions

K.K.T. and S.V.S. conceived the project. S.V.S developed the algorithm and software to analyze the data. M.K. analyzed data. K.K.T., Y.H., J.W.K.H., and S.V.S. wrote the paper. All authors commented on and edited the text.

## Ethics declarations

### Competing interests

The authors declare no competing interests.

## Notes

https://pyvia.readthedocs.io/en/latest/

https://github.com/ShobiStassen/VIA

### Competing Interest Statement

The authors have declared no competing interest.

## References

1. Sikkema, L., Ramírez-Suástegui, C., Strobl, D.C. et al. An integrated cell atlas of the lung in health and disease. Nat Med 29, 1563–1577 (2023). 10.1038/s41591-023-02327-2

2. Qiu, C., Cao, J., Martin, B.K. et al. Systematic reconstruction of cellular trajectories across mouse embryogenesis. Nat Genet 54, 328–341 (2022). 10.1038/s41588-022-01018-x

3. Lange, Merlin et al., Zebrahub – Multimodal Zebrafish Developmental Atlas Reveals the State Transition Dynamics of Late Vertebrate Pluripotent Axial Progenitors Preprint at bioRxiv 2023.03.06.531398; doi: https://doi.org/10.1101/2023.03.06.531398 (2023)

4. Grover A, Leskovec J. node2vec: Scalable Feature Learning for Networks. KDD. 2016 Aug;2016:855–864. doi: 10.1145/2939672.2939754. PMID: 27853626; PMCID: PMC5108654.

5. Renming Liu, Arjun Krishnan, PecanPy: a fast, efficient and parallelized Python implementation of node2vec, Bioinformatics, Volume 37, Issue 19, October 2021, Pages 3377–3379, 10.1093/bioinformatics/btab202

6. Renming Liu and others, Accurately modeling biased random walks on weighted networks using node2vec+, Bioinformatics, Volume 39, Issue 1, January 2023, btad047, 10.1093/bioinformatics/btad047

7. Moon, K.R., van Dijk, D., Wang, Z. et al. Visualizing structure and transitions in high-dimensional biological data. Nat Biotechnol 37, 1482–1492 (2019). 10.1038/s41587-019-0336-3

8. Lange, M., Bergen, V., Klein, M. et al. CellRank for directed single-cell fate mapping. Nat Methods 19, 159–170 (2022). 10.1038/s41592-021-01346-6

9. Pijuan-Sala, B., Griffiths, J.A., Guibentif, C. et al. A single-cell molecular map of mouse gastrulation and early organogenesis. Nature 566, 490–495 (2019). 10.1038/s41586-019-0933-9

10. Lange L, Morgan M, Schambach A. The hemogenic endothelium: a critical source for the generation of PSC-derived hematopoietic stem and progenitor cells. Cell Mol Life Sci. 2021 May;78(9):4143–4160. doi: 10.1007/s00018-021-03777-y. Epub 2021 Feb 9. PMID: tham33559689; PMCID: PMC8164610.

11. Wolf, F.A., Hamey, F.K., Plass, M. et al. PAGA: graph abstraction reconciles clustering with trajectory inference through a topology preserving map of single cells. Genome Biol 20, 59 (2019). 10.1186/s13059-019-1663-x

12. Bergen, V., Lange, M., Peidli, S. et al. Generalizing RNA velocity to transient cell states through dynamical modeling. Nat Biotechnol 38, 1408–1414 (2020). 10.1038/s41587-020-0591-3

13. Edri S, Hayward P, Jawaid W, Martinez Arias A. Neuro-mesodermal progenitors (NMPs): a comparative study between pluripotent stem cells and embryo-derived populations. Development.; 146(12):dev180190. doi: 10.1242/dev.180190. PMID: 31152001; PMCID: PMC6602346. (2019)

14. Li, M.M., Huang, K. & Zitnik, M. Graph representation learning in biomedicine and healthcare. Nat. Biomed. Eng 6, 1353–1369 (2022). 10.1038/s41551-022-00942-x

15. Canu, G., Ruhrberg, C. First blood: the endothelial origins of hematopoietic progenitors. Angiogenesis 24, 199–211 (2021). 10.1007/s10456-021-09783-9

16. Hayashi M, Pluchinotta M, Momiyama A, Tanaka Y, Nishikawa S, Kataoka H. Endothelialization and altered hematopoiesis by persistent Etv2 expression in mice. Exp Hematol. 2012 Sep;40(9):738–750.e11. doi: 10.1016/j.exphem.2012.05.012. Epub 2012 Jun 1. PMID: 22659386. (2012)

17. Jun Shen et al., Single-cell transcriptome of early hematopoiesis guides arterial endothelial-enhanced functional T cell generation from human PSCs.Sci. Adv.7,eabi9787(2021).DOI:10.1126/sciadv.abi9787 (2021)

18. Gao L., Tober J., Gao P., Chen C., Tan K. and Speck N. A. (2018). RUNX1 and the endothelial origin of blood. Exp. Hematol. 68, 2–9. 10.1016/j.exphem.2018.10.009

19. Thambyrajah R., Mazan M., Patel R., Moignard V., Stefanska M., Marinopoulou E., Li Y., Lancrin C., Clapes T., Möröy T. et al. (2016a). GFI1 proteins orchestrate the emergence of haematopoietic stem cells through recruitment of LSD1. Nat. Cell Biol. 18, 21–32. 10.1038/ncb3276

20. Cambray N, Wilson V. Two distinct sources for a population of maturing axial progenitors. Development. 2007 Aug;134(15):2829–40. doi: 10.1242/dev.02877. Epub 2007 Jul 4. PMID: 17611225

21. Duan D, Fu Y, Paxinos G, Watson C. Spatiotemporal expression patterns of Pax6 in the brain of embryonic, newborn, and adult mice. Brain Struct Funct. 2013 Mar;218(2):353–72. doi: 10.1007/s00429-012-0397-2. Epub 2012 Feb 22. PMID: 22354470

22. Haedicke J, Brown C, Naghavi MH. The brain-specific factor FEZ1 is a determinant of neuronal susceptibility to HIV-1 infection. Proc Natl Acad Sci U S A. 2009 Aug 18;106(33):14040–5. doi: 10.1073/pnas.0900502106. Epub 2009 Aug 10. PMID: 19667186; PMCID: PMC2729016

23. Cheung M, Briscoe J. Neural crest development is regulated by the transcription factor Sox9. Development. 2003 Dec;130(23):5681–93. doi: 10.1242/dev.00808. Epub 2003 Oct 1. PMID: 14522876.

24. Mork L, Crump G. Zebrafish Craniofacial Development: A Window into Early Patterning. Curr Top Dev Biol. 2015;115:235–69. doi: 10.1016/bs.ctdb.2015.07.001. Epub 2015 Oct 6. PMID: 26589928; PMCID: PMC4758817.

25. Knight RD, Schilling TF. Cranial neural crest and development of the head skeleton. Adv Exp Med Biol. 2006;589:120–33. doi: 10.1007/978-0-387-46954-6_7. PMID: 17076278.

26. Steventon B, Mayor R, Streit A. Neural crest and placode interaction during the development of the cranial sensory system. Dev Biol. 2014 May 1;389(1):28–38. doi: 10.1016/j.ydbio.2014.01.021. Epub 2014 Jan 31. PMID: 24491819; PMCID: PMC4439187

27. Steventon B. and Martinez Arias A. (2017). Evo-engineering and the cellular and molecular origins of the vertebrate spinal cord. Dev. Biol. 432, 3–13. 10.1016/j.ydbio.2017.01.021

28. Chatterjee M, Li JY. Patterning and compartment formation in the diencephalon. Front Neurosci. 2012 May 11;6:66. doi: 10.3389/fnins.2012.00066. PMID: 22593732; PMCID: PMC3349951

29. Ohyama T, Mohamed OA, Taketo MM, Dufort D, Groves AK. Wnt signals mediate a fate decision between otic placode and epidermis. Development. 2006 Mar;133(5):865–75. doi: 10.1242/dev.02271. Epub 2006 Feb 1. PMID: 16452098.

30. Maier EC, Saxena A, Alsina B, Bronner ME, Whitfield TT. Sensational placodes: neurogenesis in the otic and olfactory systems. Dev Biol. 2014 May 1;389(1):50–67. doi: 10.1016/j.ydbio.2014.01.023. Epub 2014 Feb 6. PMID: 24508480; PMCID: PMC3988839

31. Teixeira Rosa J, Oralová V, Larionova D, Eisenhoffer GT, Eckhard Witten P, Huysseune A. Periderm invasion contributes to epithelial formation in the teleost pharynx. Sci Rep. 2019 Jul 12;9(1):10082. doi: 10.1038/s41598-019-46040-y. PMID: 31300674; PMCID: PMC6626026.

32. van der Maaten, L. & Hinton, G. (2008), ‘Visualizing Data using t-SNE ‘, Journal of Machine Learning Research 9, 2579--2605.

33. Wymeersch FJ, Skylaki S, Huang Y, Watson JA, Economou C, Marek-Johnston C, Tomlinson SR, Wilson V. Transcriptionally dynamic progenitor populations organised around a stable niche drive axial patterning. Development. 2019 Jan 2;146(1):dev168161. doi: 10.1242/dev.168161. PMID: 30559277; PMCID: PMC6340148.

34. Nowotschin S, Setty M, Kuo YY, Liu V, Garg V, Sharma R, Simon CS, Saiz N, Gardner R, Boutet SC, Church DM, Hoodless PA, Hadjantonakis AK, Pe’er D. The emergent landscape of the mouse gut endoderm at single-cell resolution. Nature. 2019 May;569(7756):361-367. doi: 10.1038/s41586-019-1127-1. Epub 2019 Apr 8. PMID: 30959515; PMCID: PMC6724221.

35. Balmer S, Nowotschin S, Hadjantonakis AK. Notochord morphogenesis in mice: Current understanding & open questions. Dev Dyn. 2016 May;245(5):547–57. doi: 10.1002/dvdy.24392. Epub 2016 Mar 14. PMID: 26845388; PMCID: PMC4844759

36. Kwon GS, Viotti M, Hadjantonakis AK. The endoderm of the mouse embryo arises by dynamic widespread intercalation of embryonic and extraembryonic lineages. Dev Cell. 2008 Oct;15(4):509–20. doi: 10.1016/j.devcel.2008.07.017. PMID: 18854136; PMCID: PMC2677989.

37. Henrique D, Abranches E, Verrier L, Storey KG. Neuromesodermal progenitors and the making of the spinal cord. Development. 2015 Sep 1;142(17):2864–75. doi: 10.1242/dev.119768. PMID: 26329597; PMCID: PMC4958456.

38. Wilson V., Olivera-Martinez I. and Storey K. G. (2009). Stem cells, signals and vertebrate body axis extension. Development 136, 1591–1604. 10.1242/dev.021246

39. Aitana Perea-Gomez and Sigolene M. Meilhac, Formation of the Anterior-Posterior Axis in Mammals, https://api.semanticscholar.org/CorpusID:80823225 (2015)

40. Li H, Chang YW, Mohan K, Su HW, Ricupero CL, Baridi A, Hart RP, Grumet M. Activated Notch1 maintains the phenotype of radial glial cells and promotes their adhesion to laminin by upregulating nidogen. Glia. 2008 Apr 15;56(6):646–58. doi: 10.1002/glia.20643. PMID: 18286610; PMCID: PMC2712347.

41. Dang L, Yoon K, Wang M, Gaiano N. Notch3 signaling promotes radial glial/progenitor character in the mammalian telencephalon. Dev Neurosci. 2006;28(1-2):58–69. doi: 10.1159/000090753. PMID: 16508304.

42. Dimou L, Götz M. Glial cells as progenitors and stem cells: new roles in the healthy and diseased brain. Physiol Rev. 2014 Jul;94(3):709–37. doi: 10.1152/physrev.00036.2013. PMID: 24987003.

43. Wang K, Hou L, Wang X, Zhai X, Lu Z, Zi Z, Zhai W, He X, Curtis C, Zhou D, Hu Z. PhyloVelo enhances transcriptomic velocity field mapping using monotonically expressed genes. Nat Biotechnol. 2023 Jul 31. doi: 10.1038/s41587-023-01887-5. Epub ahead of print. PMID: 37524958.

44. Christophe Hurter, Ozan Ersoy, Alexandru C Telea. Graph bundling by Kernel Density Estimation. EUROVIS 2012, Eurographics Conference on Visualization, Jun 2012, Vienna, Austria. pp 865–874, ff10.1111/j.1467-8659.2012.03079.xff. Ffhal-01022472f

45. Quake SR. A decade of molecular cell atlases. Trends Genet. 2022 Aug;38(8):805–810. doi: 10.1016/j.tig.2022.01.004. Epub 2022 Jan 31. PMID: 35105475.

46. Calderon D, Blecher-Gonen R, Huang X, Secchia S, Kentro J, Daza RM, Martin B, Dulja A, Schaub C, Trapnell C, Larschan E, O’Connor-Giles KM, Furlong EEM, Shendure J. The continuum of Drosophila embryonic development at single-cell resolution. Science. 2022 Aug 5;377(6606):eabn5800. doi: 10.1126/science.abn5800. Epub 2022 Aug 5. PMID: 35926038; PMCID: PMC9371440.

47. The Tabula Sapiens Consortium*,The Tabula Sapiens: A multiple-organ, single-cell transcriptomic atlas of humans.Science376,eabl4896(2022).DOI:10.1126/science.abl4896

48. Shobana V Stassen, Dickson M D Siu, Kelvin C M Lee, Joshua W K Ho, Hayden K H So, Kevin K Tsia, PARC: ultrafast and accurate clustering of phenotypic data of millions of single cells, Bioinformatics, Volume 36, Issue 9, May 2020, Pages 2778–2786, 10.1093/bioinformatics/btaa042

49. Rajewsky, N., Almouzni, G., Gorski, S.A. et al. LifeTime and improving European healthcare through cell-based interceptive medicine. Nature 587, 377–386 (2020). 10.1038/s41586-020-2715-9

50. McInnes, L., Healy, J., Saul, N. & Großberger, L. UMAP: uniform manifold approximation and projection. J. Open Source Softw. 3, 861 (2018).

51. Setty, M., Kiseliovas, V., Levine, J. et al. Characterization of cell fate probabilities in single-cell data with Palantir. Nat Biotechnol 37, 451–460 (2019). 10.1038/s41587-019-00

52. Jacomy M, Venturini T, Heymann S, Bastian M. ForceAtlas2, a continuous graph layout algorithm for handy network visualization designed for the Gephi software. PLoS One. 2014 June

53. Joseph A. Cottam, Andrew Lumsdaine, and Peter Wang “Abstract rendering: out-of-core rendering for information visualization”, Proc. SPIE 9017, Visualization and Data Analysis 2014, 90170K (3 February 2014); 10.1117/12.2041200

54. Benjamin Jurek and Inga D. Neumann, The Oxytocin Receptor: From Intracellular Signaling to Behavior, Physiological Reviews 2018 98:3, 1805–190

55. Qiu C, Martin BK, Welsh IC, et al., A single-cell transcriptional timelapse of mouse embryonic development, from gastrula to pup. bioRxiv [Preprint]. 2023 Apr 5:2023.04.05.535726. doi: 10.1101/2023.04.05.535726. PMID: 37066300; PMCID: PMC10104014.

56. George Paxinos, Keith B.J. Franklin, Paxinos and Franklin’s the Mouse Brain in Stereotaxic Coordinates, 5th Edition 2019, Academic Press, eBook ISBN: 9780128161586

