## Supplementary Information for "StaVia: Spatially and temporally aware cartography with higher order random walks for cell atlases"

### Supplementary Materials:

#### List of Figures

Supplementary Fig.S1. Comparing TI graph structure of Via2.0 and CellRank+PAGA
Supplementary Fig.S2. Lineage paths and gene trends for cell fates in Via 2.0 using memory
Supplementary Fig.S3. Lineage paths and gene trends comparison between Via 2.0 and CellRank
Supplementary Fig.S4. Via 2.0 Memory helps distinguish gene trends near the NMP lineages
Supplementary Fig.S5. Stability analysis for memory
Supplementary Fig.S6. CellRank lineage probabilities not improved by projecting on Atlas View
Supplementary Fig.S7. Zebrahub mesoderm: Comparison of lineage paths and gene specificity trends
Supplementary Fig.S8. Zebrahub neural ectoderm: Comparison of lineage paths and gene trends
Supplementary Fig.S9. Zebrahub non-neural ectoderm: Comparison of lineage paths and gene trends.
Supplementary Fig S10. Impact of Memory on Mesodermal Lineage Probabilities
Supplementary Fig S11. Impact of Memory on Mesodermal Gene Trends (Zebrahub)
Supplementary Fig.S12 Comparison of visualization methods on different time-series RNA-seq datasets (cells colored by tissue type)
Supplementary Fig.S13 Comparison of visualization methods on different time-series RNA-seq datasets (cells colored by developmental stage)
Supplementary Fig.S14 Impact of key steps in Via 2.0 Atlas View embedding
Supplementary Fig.S15 Impact of Steps in Via 2.0 Embedding (cells colored by developmental stage)
Supplementary Fig.S16 Impact of Steps in Via 2.0 Embedding (cells colored by tissue type)
Supplementary Fig.S17 Impact of Steps in Via 2.0 Embedding (Radar plots)

#### List of Notes and Tables

Supplementary Note 1: Selection of TI methods used for comparison
Supplementary Note 2: Via reveals 3 developmental patterns in Murine Embryogenesis
Supplementary Table 1 for parameters used in TI
Supplementary Table 2 for parameters used for single-cell embeddings

### Supplementary Note 1: Selection of TI methods used for comparison

We focus on comparing Via 2.0 to a hybrid pipeline of CellRank+PAGA+scVelo as this allows for a more apples-to-apples comparison with some key features in Via 2.0 such as:

1. Combining feature-distances, pseudotime and RNA velocity information to infer direction such that datasets without RNA-velocity (or where velocity is too noisy to be used exclusively) can still be analyzed using CellRank and PAGA. We note that the pseudotime-transition matrix in CellRank is computed using either Via 1.0 or Palantir. Incorporating the temporal annotations in time-series data is not available in CellRank.
2. Interchangeable interpretations of TI and visualization at both the cluster graph level and single-cell level lineages (pseudotimes, cell fates and lineage probabilities) using the PAGA-CellRank cell-fate cluster graphs (**Fig.S1 iv and Fig. S6c** “cell fate view”) and scVelo-directed-PAGA graphs alongside, scVelo-CellRank gene trends and CellRank’s single-cell lineage probabilities projected onto a UMAP/t-SNE.

In addition to CellRank, scVelo and directed-Paga, there are a handful of other RNA-velocity based TI methods that have emerged in the past two years. These are Cytopath [Gupta 2022], Cellpath [Z. Zhang and X.Zhang 2021] and Vetra [G.Weng 2021].

With respect to our first criteria of offering a ‘hybrid’ approach that allows integration of non-RNA velocity based and velocity based direction inference, we note that these three methods rely exclusively on RNA-velocity to infer direction and are limited to use on small datasets of less than 10,000 cells. Cytopath relies on scVelo’s pipeline (or the user) to provide the transition probability matrix, the root and terminal states, three elements which can substantially influence accuracy of TI analysis. Because of their reliance on the quality of the RNA-velocity data, Vetra and Cellpath often suffer from poor root and terminal state detection [Gupta et al., 2022] and cannot accept user-defined roots/cell fates to provide adjustments, which can distort results. Cellpath also frequently grossly overestimates the number of trajectories which confounds downstream analysis [Gupta et al., 2022] and requires manual subselection.

With respect to our second criteria, we note that the methods described above do not offer an intuitive way to liaise visually or computationally between the cluster graph and single-cell resolution trajectory which is a unique aspect of Via 2.0’s output. However, combining Paga+scVelo+CellRank allows us to conduct a more apples-to-apples comparison. In our analysis, this generally required manual assignment in CellRank of several cell fates that were detected by Via 2.0. Via 2.0 completes the full trajectory computation in 3 minutes, compared to CellRank’s 20 minutes during which 8 Intel Zeon 3.6GHz CPUs are fully occupied. Pre-processing and sc-embedding computation runtime are excluded in these numbers. The computation time of scVelo’s latent time which is used in CellRank’s gene trends is also excluded from the reported runtime and takes (on the Mouse Gastrulation dataset) 105 minutes to compute on 8 cores and requires 99 GB RAM, compared to Via 2.0’s computes in ~ 1 minute and requires less than half the peak RAM.

### Supplementary Note 2: Via 2.0 reveals developmental structures using its directed graph layout

Here we highlight three novel developmental transitions (with reference to **Fig.2 and Fig.3a in Results**) not automatically captured in other TI methods nor clearly visualized in other representations of the full dataset.

**1. Hematopoietic waves:** The Atlas View and the cluster graph reveal two waves of hematopoiesis - primitive and pro-definitive. The spatial arrangement of cells on these graphs in terms of adjacent populations and temporal stage is consistent with knowledge that the transient primitive (first) wave arises shortly after mesoderm formation [L. Lange 2021, Palis 2014]. In Via 2.0, the hematopoietic potential of the pro-definitive (second wave) is marked by an E8.5 population of hemogenic endothelial cells which undergo an endothelial-to-hematopoietic transition (EHT)[Canu & Ruhrberg 2021]. Amongst the hemato-endothelial *Kdr+Cd34+* progenitors **(Fig. 2b)**, the cells with lower *Etv2* (a hematopoiesis suppressor) are perhaps those most poised for EHT [Hayashi 2012, Shen 2021].

**2. NMPs at the neuro-mesodermal junction:** The Atlas View (**Fig.2a,c**) depicts NMPs, bipotent progenitors expressing *T (Brachyury),* that contribute to spinal cord and paraxial mesoderm during axial elongation, arising from the caudal epiblast at E8.0 [Edri 2019, Cambray and Wilson, 2007; Wymeersch 2019]. NMPs closer to the paraxial mesoderm, residing on the upper section of the triangle marked in **Fig.2c**, express higher *Tbx6* expression, suggesting a pro-mesodermal tendency, consistent with the paraxial mesoderm cells neighboring this edge of the NMP-triangle. NMPs on the lower edge of the triangle express more *Nkx1-2*, correctly indicating a propensity towards the spinal cord neural cells that lie adjacent to them on the Atlas [Henrique 2015; Steventon & Arias 2017, Wilson 2009, Edri 2019]. **Fig.3d** shows that the sequential gene expression of gastrulation to NMP is automatically captured along the pseudotime axis for the NMP lineage.

**3. Gut development:** In contrast to prior analysis [Pijuan-Sala 2019] which required subsetting of the visceral and definitive endoderm (VE, DE) and gut cells *(Wnt5b+)* to observe the dual origin of the gut, Via 2.0's cluster graph **(Fig.3a)** and Atlas View **(Fig. 2a,d)** reveal that gut endoderm morphogenesis arises from two distinct developmental origins: the intercalation of the streak derived *Sox17* positive definitive endoderm (DE) and the dispersal of *Ttr* expressing extraembryonic VE [Kwon 2008, Balmer, 2017, Nowotschin 2019].

### Supplementary Tables for parameters used in TI

#### Table S1.a. Mouse Gastrulation:

| **Method** | **Common** | **Unique Parameters** |
|---|---|---|
| Via 2.0 | PCs = 30<br>KNN =30 | Velo_weight = 0.5<br>Knn_seq = 10 (20knn +10knnseq = 30 knn total) |
| CellRank | | N_macrostates = 15 (default = 10, but increasing to 15 improved results). Higher values than 15 took prohibitively long runtimes without improvement |
| PAGA | | scVelo's latent time for direction (uses CellRank's initial and terminal states) |

#### Table S1.b. Zebrahub:

| **Method** | **Common** | **Unique Parameters** |
|---|---|---|
| Via 2.0 | PCs = 100<br>KNN =15 | Velo_weight = 0.5<br>Num_seq_knn = 5 (i.e. 10 non-seq, and 5seq = 15 knn) |
| CellRank | | N_macrostates = 15 |
| PAGA | | Direction based on scVelo's latent time (or diffusion pseudotime for which root state was provided) |

### Supplementary Tables for parameters used in visualization

All methods are run on default parameters. Non-default parameters are listed below.

#### Table S2.

| Dataset | Common | Via 2.0 | Umap | t-SNE |
|---|---|---|---|---|
| Mouse Gastrulation | PCs = 30<br>KNN =30 | min_dist=0.2<br>30knn=15seq +15 nonseq | min_dist=0.2 | Perplexity =30 |
| Zebrahub Lange | PCs = 100<br>KNN =30 | min_dist=0.3<br>30knn=15seq +15 nonseq | min_dist=0.3<br>knn=50 to reduce fragmentation | Perplexity =30 |
| Zebrafish Wagner | PCs = 30<br>knn=60 | min_dist=0.1<br>60knn=45seq +15 nonseq | min_dist=0.1 | Perplexity = 100 |
| Ascidian protovert Cao | PCs = 30<br>KNN =30 | min_dist=0.3<br>30knn=15seq +15 nonseq | min_dist=0.3 | Perplexity = 30 |
| Mouse Neuron Manno | PCs=30<br>knn=30 | min_dist=0.2<br>30knn=15seq +15 nonseq | min_dist=0.2 | Perplexity =30 |

**Supplementary Fig.S1 Comparing TI graph structure of Via2.0 and Cellrank+PAGA**

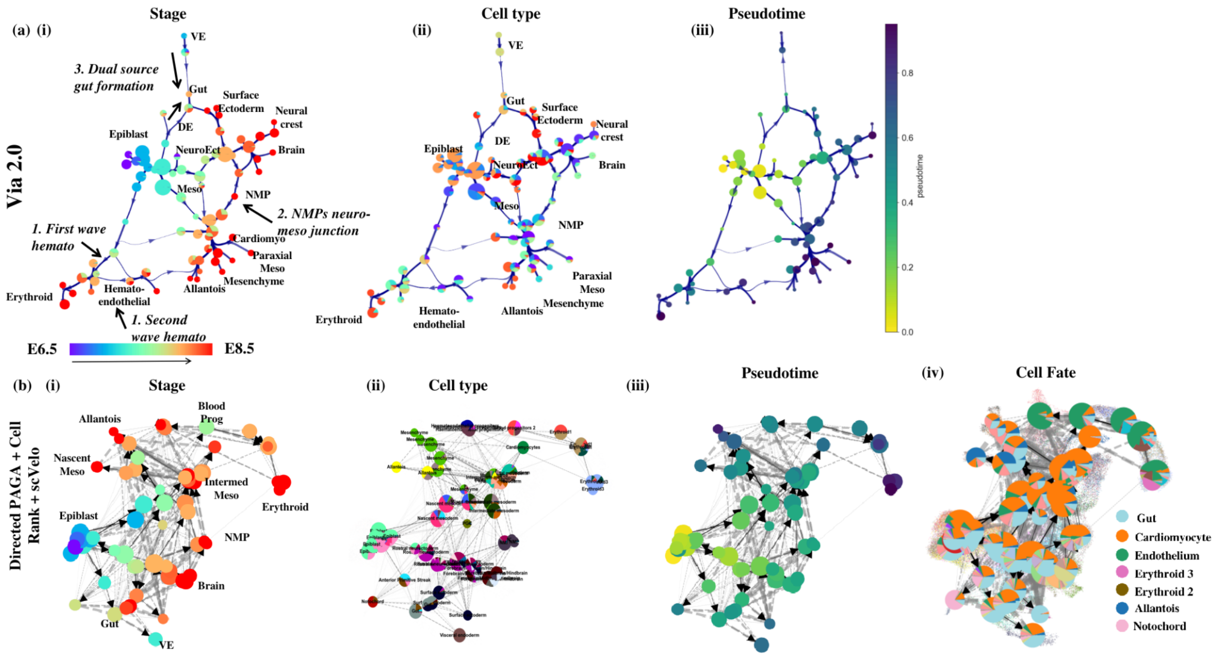


***Supplementary Fig.S1. Comparing TI graph structure of Via2.0 and CellRank+PAGA*** *(a) Via 2.0 pseudotime-RNA velocity directed cluster graph colored by (i) developmental stage (ii) cell type (iii) Via pseudotime (b) scVelo-latent time directed PAGA graph, initialized with CellRank's initial states. Gray arrows denote connectivity and black arrow-edges denote direction, colored by (i) known developmental stage (ii) cell type (iii) Pseudotime (iv) CellRank's lineage probability of cells in that cluster towards one of the detected terminal states. Due to most lineage probabilities being very localized, and a few being very diffuse over all cells (cardiomyocyte and surface ectoderm whose sc-likelihoods are shown in Fig.3 and Fig. S3 ), we see the cell fates associated with the diffused lineage probabilities over-represented in almost all clusters regardless of relevance to the cell fate.*

**Supplementary Fig.S2 Lineage paths and gene trends for cell fates in Via2.0 using memory**

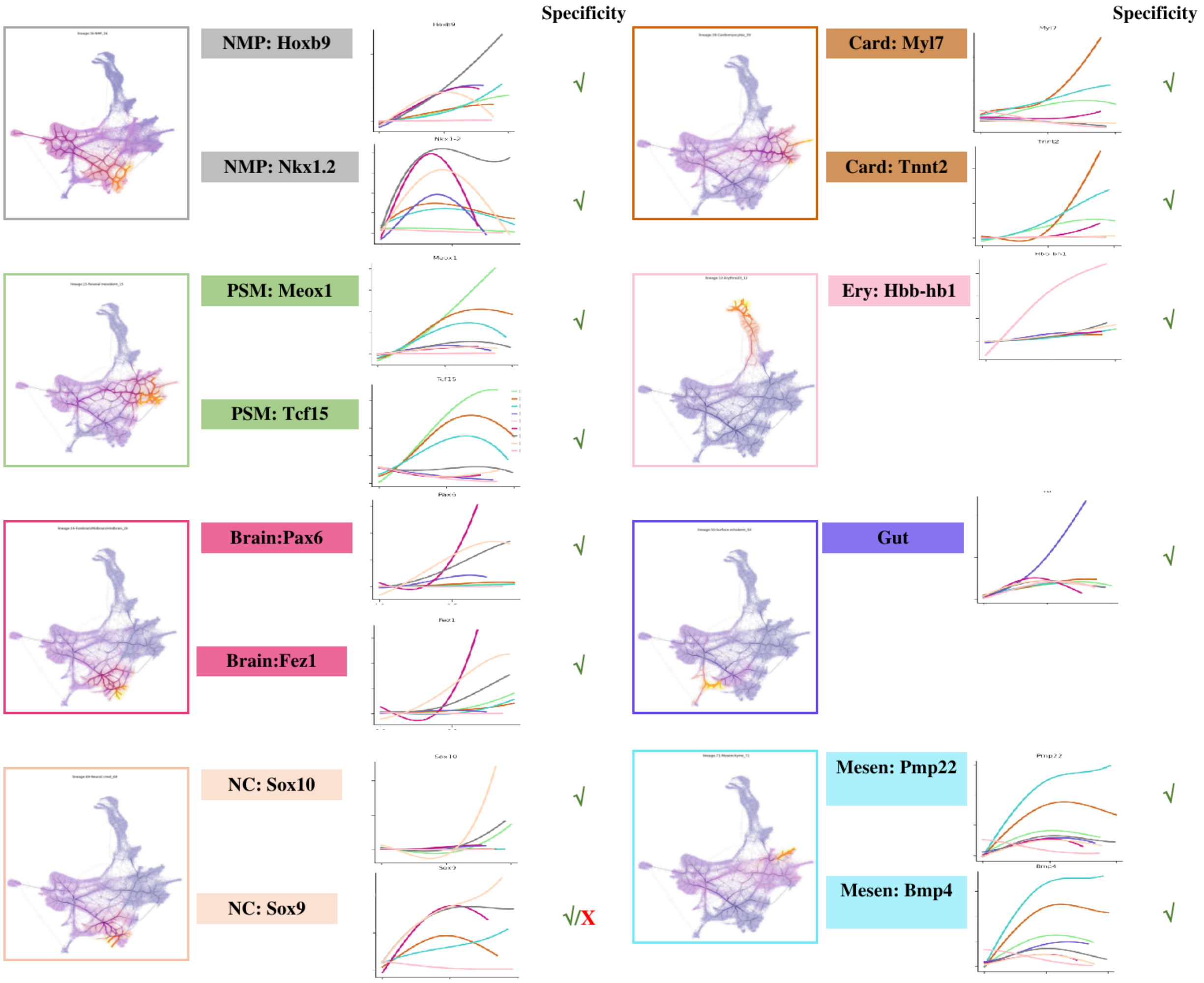


***Supplementary Fig.S2. Lineage paths and gene trends for cell fates in Via 2.0 using memory:*** *Lineage pathways detected by Via 2.0 for cell fates show progression from epiblast (E6.5) through relevant intermediate stages before arriving at the cell fate population (E8.5). Two marker genes are shown for each lineage. Each plot shows the gene expression versus pseudotime for all lineages. The lineage trend of interest is indicated by the color of the gene-box and the border of the lineage plot. If the correct lineage shows upregulation of the highlighted marker gene, then the color of the trend line and gene label/plot border will match. E.g. The cardiomyocyte lineage is light brown. In the gene trends for Myl7 and Tnnt2, the brown cardiomyocyte trend is the single most upregulated compared to other trend lines belonging to the other lineages.*

**Supplementary Fig.S3. Comparison of Lineage paths and gene trends between Via 2.0 and Cellrank**

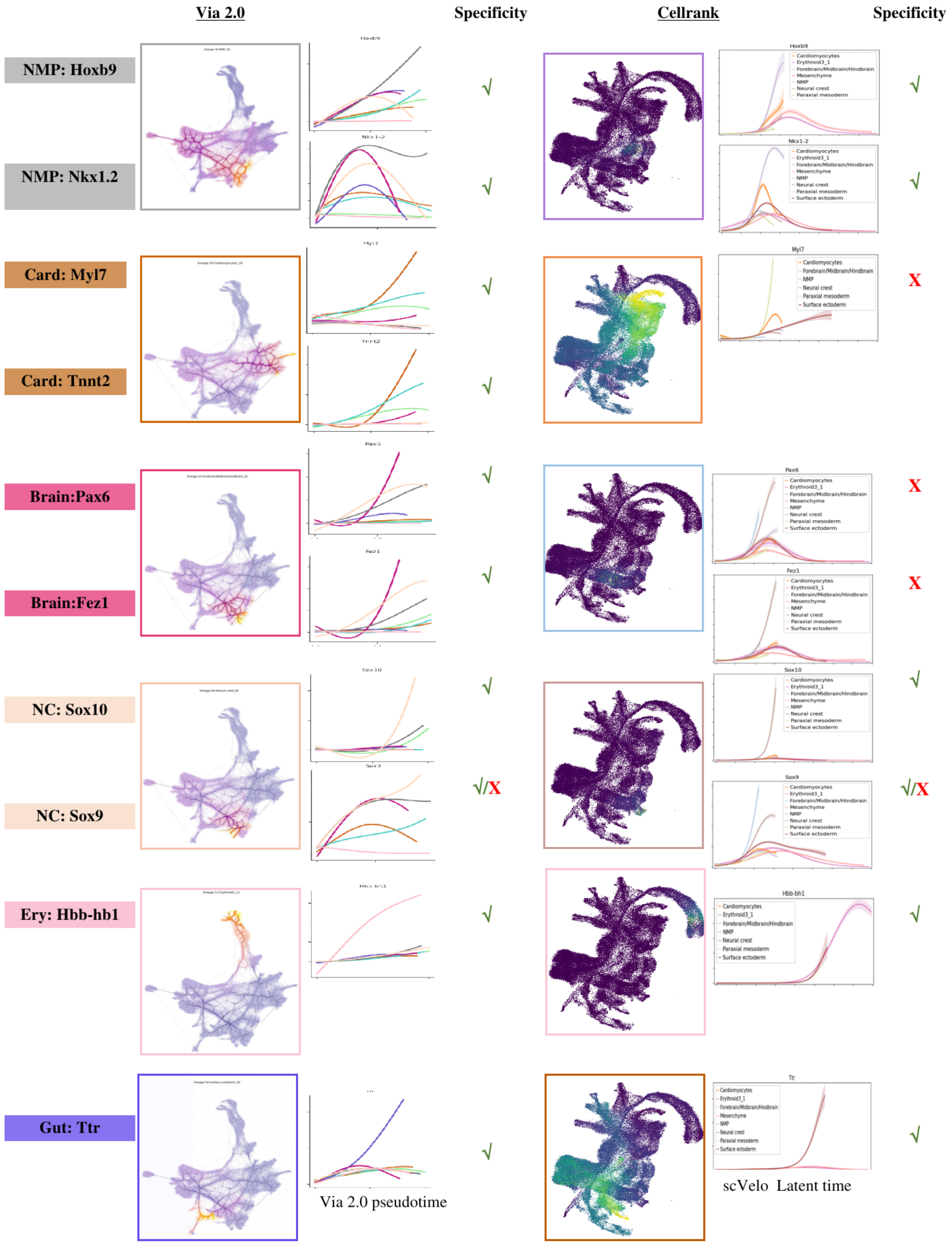


***Supplementary Fig.S3 Lineage paths and gene trends comparison between Via 2.0 and CellRank.*** *CellRank's lineage pathways are either very localized and show little information on the end-to-end progression from epiblast along consecutive stages, or are very diffuse (Gut), such that several cell populations are shown to progress towards the gut. A consequence of this is that the gene trend lines associated with lineages are not correctly upregulated. For example, the Brain lineage trend line is light blue in CellRank (indicated by the border color of the pathway plot). However, the light blue gene trend line is not the most upregulated for Pax6 or Fez1 and hence receives a cross-mark in the columns titled "specificity". scVelo's latent time used for gene trends takes 105 minutes to compute on 8 cores and requires 99 GB RAM, compared to Via 2.0's computes in ~ 1 minute and requires less than half the peak RAM.*

**Supplementary Fig.S4. Via 2.0 Memory helps distinguish gene trends near the NMP lineages**

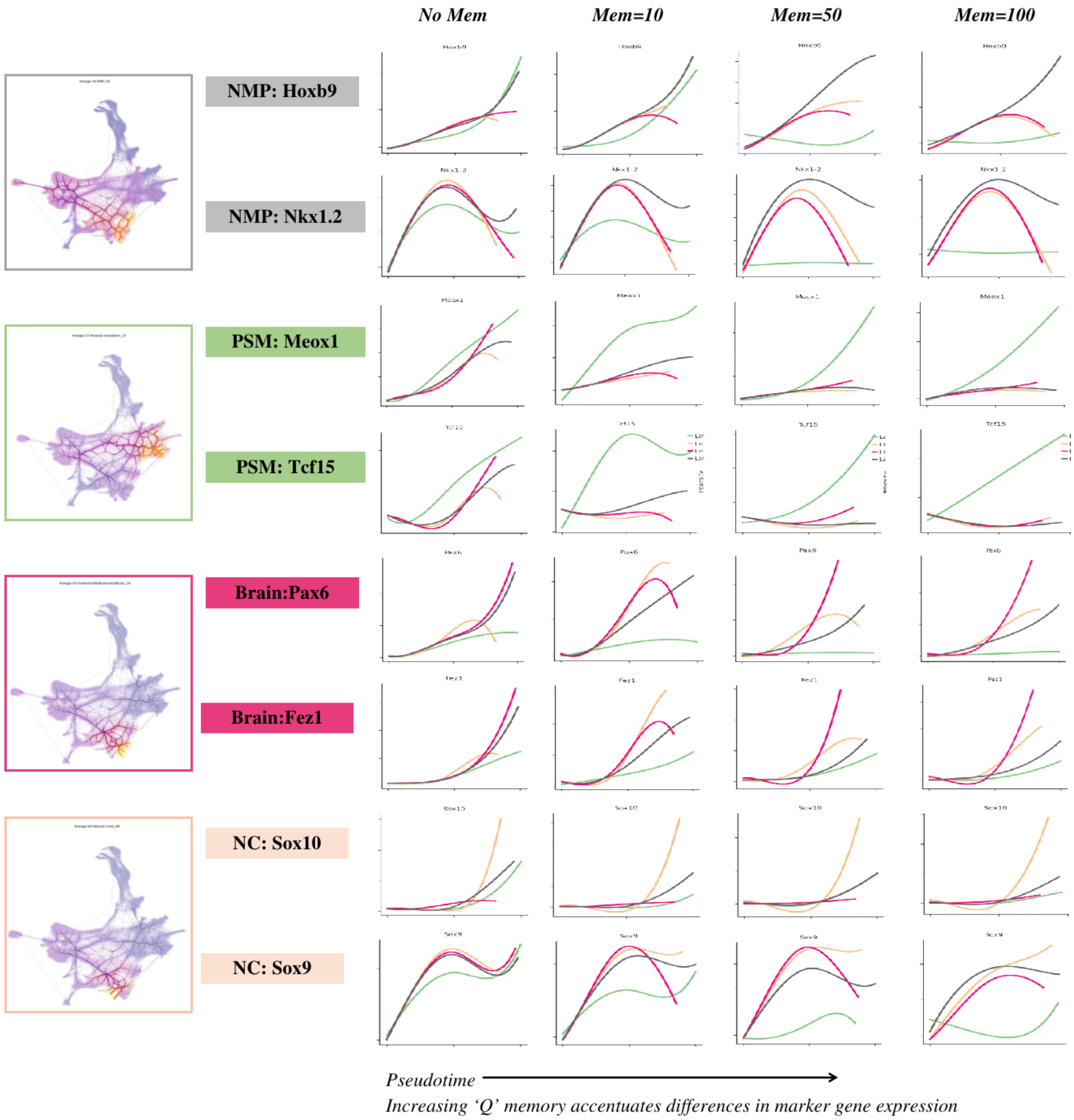


***Supplementary Fig.S4. Via 2.0 Memory helps distinguish gene trends near the NMP lineages:*** *Each gene trend subplot shows the gene expression along pseudotime for four lineages: NMP (neural mesodermal progenitors), PSM (Presomitic/paraxial mesoderm), Brain and NC (Neural crest) which are seen to emerge in varying proximity to each other. In particular with the NMPs closely linked to the brain and PSM populations, and the NC and Brain also being closely related. As a result, it can be challenging to obtain pathways that are specific enough to the relevant cell fates such that the correct gene trends for the relevant cell fate is plotted. By increasing memory, the correct lineage is upregulated for the marker gene, while other lineages remain suppressed.*

### Supplementary Fig.S5 Stability analysis for memory

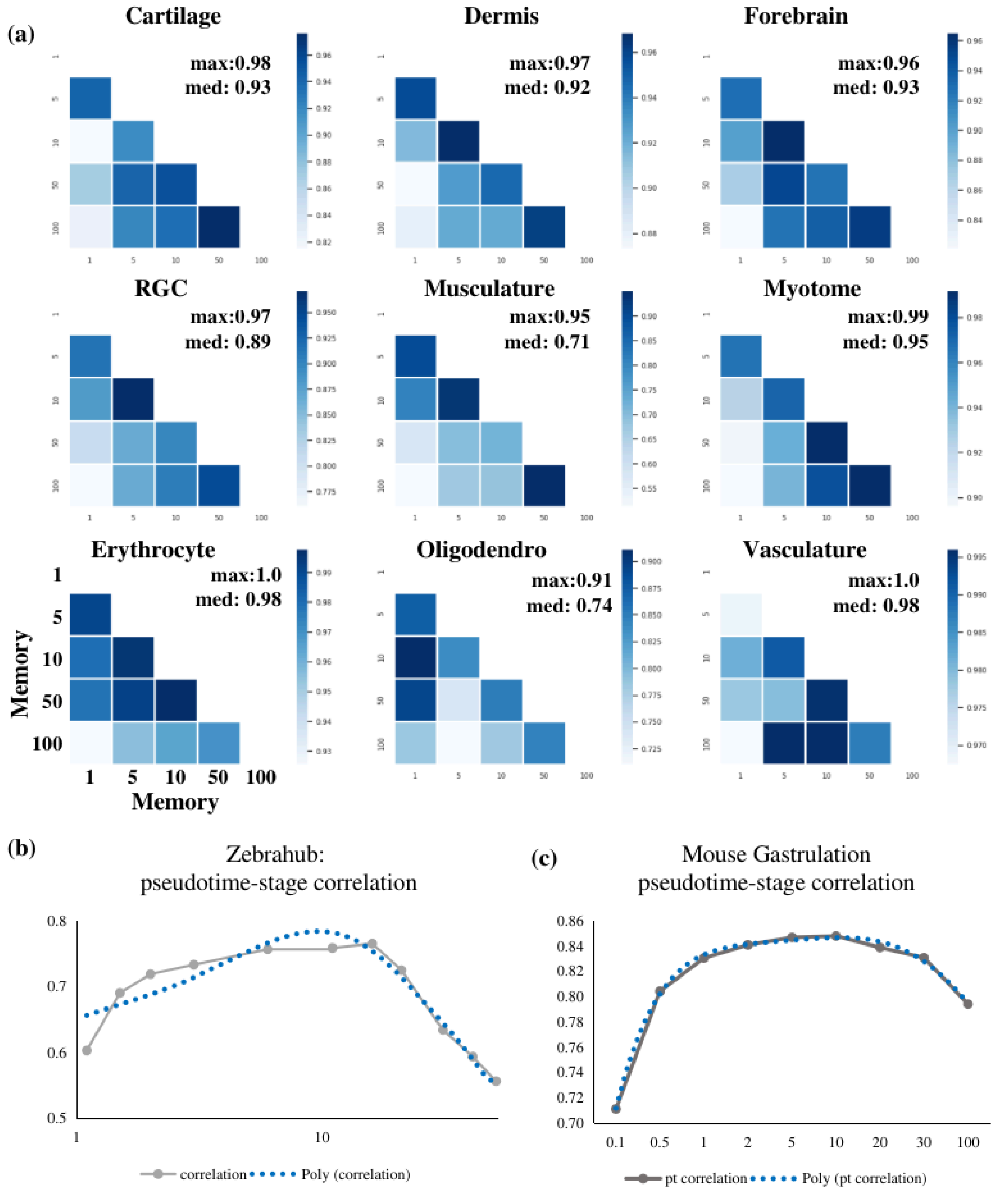


***Supplementary Fig.S5 Stability analysis for memory*** *(a) pairwise correlation of lineage probabilities for Zebrahub cell fates when increasing memory from No memory (=1) to Memory =100 (b) for memory (value of 1 signified No memory, (b) correlation of stage-labels and inferred pseudotime of memory values 0.01-50. x-axis is plotted on log(1+memory) base10 scale. Shows that values between 5-20 are desirable for Zebrahub. (c) correlation between inferred pseudotime to Mouse gastrulation stages for different memory values.*

**Supplementary Fig.S6 Zebrahub: Cellrank lineage probabilities not improved by projecting on different layout/embeddings**

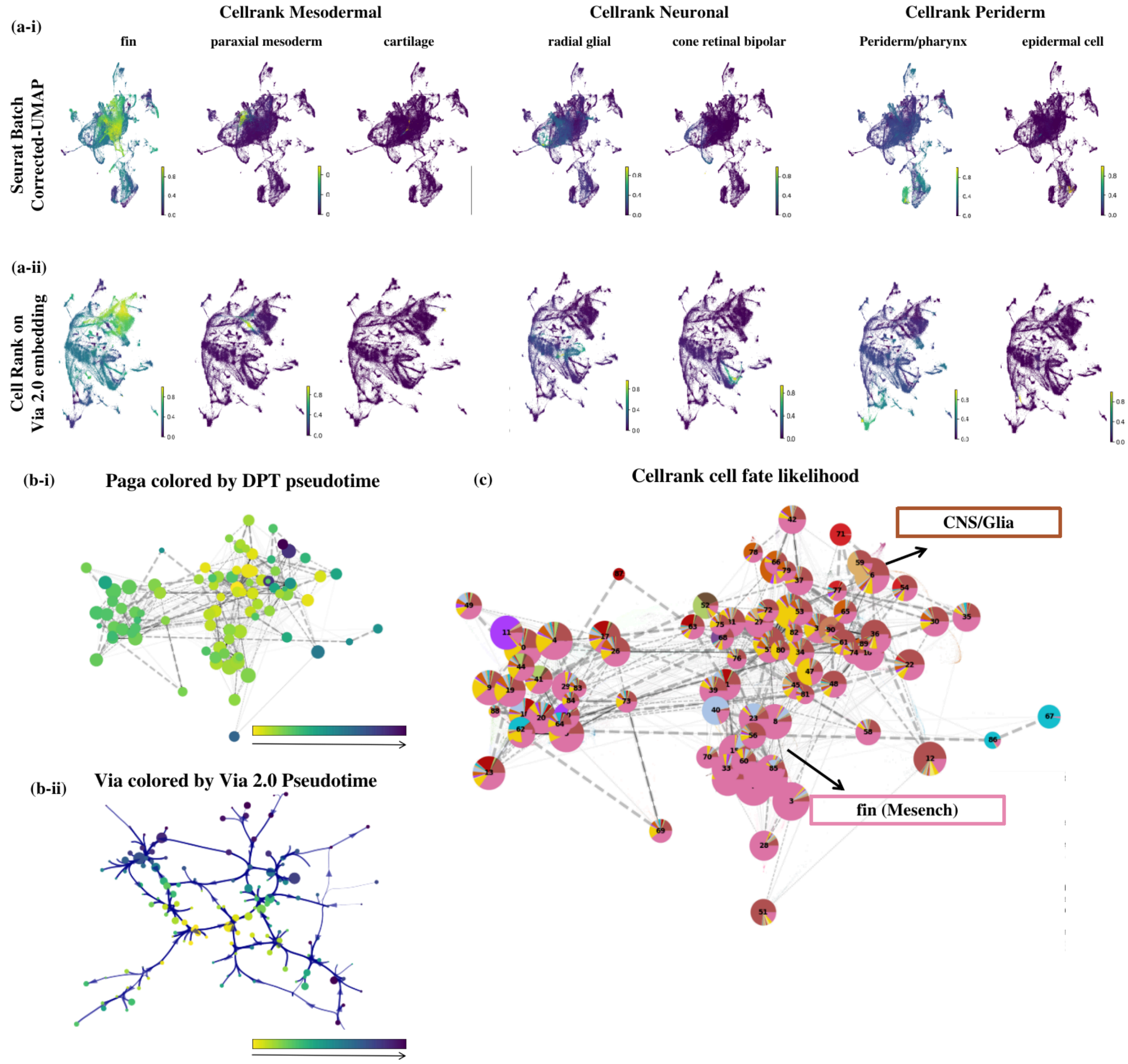


***Supplementary Fig.S6 Zebrahub: CellRank lineage probabilities does not improve by projecting on different layout/embeddings*** *(a) Automatically captured cell fates for CellRank are shown here. The lack of end-to-end information in CellRank lineage probabilities cannot be corrected by simply using a different layout for the single-cell embedding, we plot CellRank's lineage probabilities for each of its automatically detected cell fates on the UMAP on batch corrected PCs provided in the publicly available data file for Zebrahub (Anndata file without RNA velocity) as well as on Via 2.0's Atlas sc-embedding (b) Paga cluster graph colored by DPT pseudotime shows how the pseudotime scale is distorted. (c) Cluster graph composition is colored by cells' lineage probabilities towards one of the detected terminal states. Due to the highly diffused lineage probabilities shown in (a) for the fin lineage, we see that the "pink" fin terminal state is overrepresented in the CellRank-paga cell fate cluster graph. It incorrectly suggests that all cells are moving primarily towards the pink fin cell fate or the brick-brown Glia*

### Supplementary Fig.S7. Zebrahub mesoderm: Comparison of lineage paths and gene specificity trends

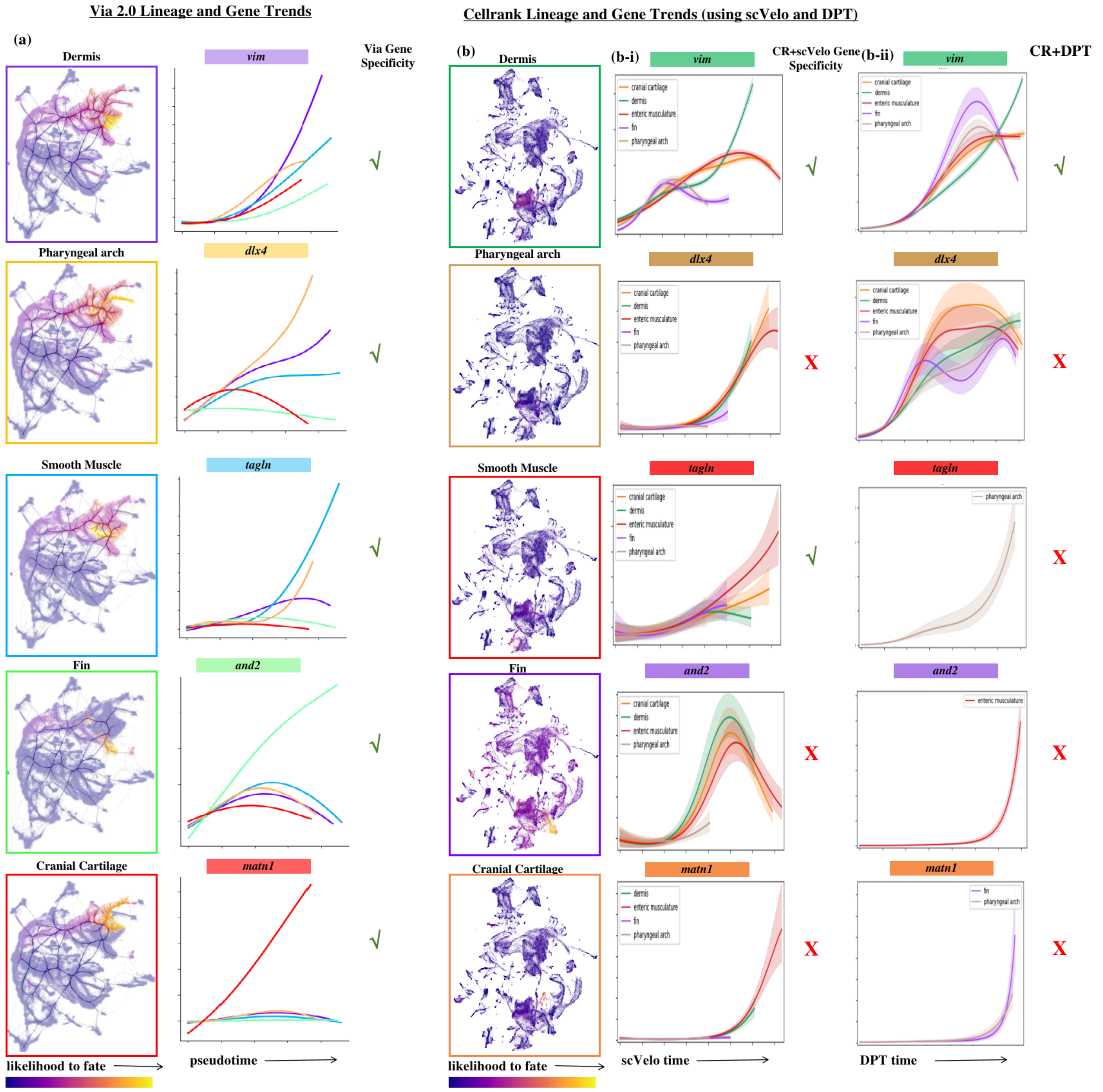


***Supplementary Fig.S7. Zebrahub mesoderm: Comparison of lineage paths and gene specificity trends*** *(a) End-to-end lineage probabilities for automatically detected mesodermal cell fates in Via 2.0 at Memory =10. All mesodermal-lineage gene trends are plotted, but only the lineage for which the marker gene is relevant, is upregulated in each case - indicated by the green tick mark which means the color of the trendline matches the color of the associated gene name. (b) In order to facilitate comparison,* ***we manually assign cell fates for CellRank which does not detect all relevant fates.*** *CellRank's pathways projected on the UMAP used in the Zebrahub paper and made publicly available (Anndata object with velocity) (b-i) shows the gene trends for the same mesodermal lineages (we manually assign those that are not detected by CellRank in order to compare all cell fate pathways) when CellRank's lineage probabilities are used together with scVelo's latent time. (b-ii) Gene trends for CellRank when using the DPT pseudotime*

**Supplementary Fig.S8. Zebrahub comparison of neuronal lineage paths and gene trends**

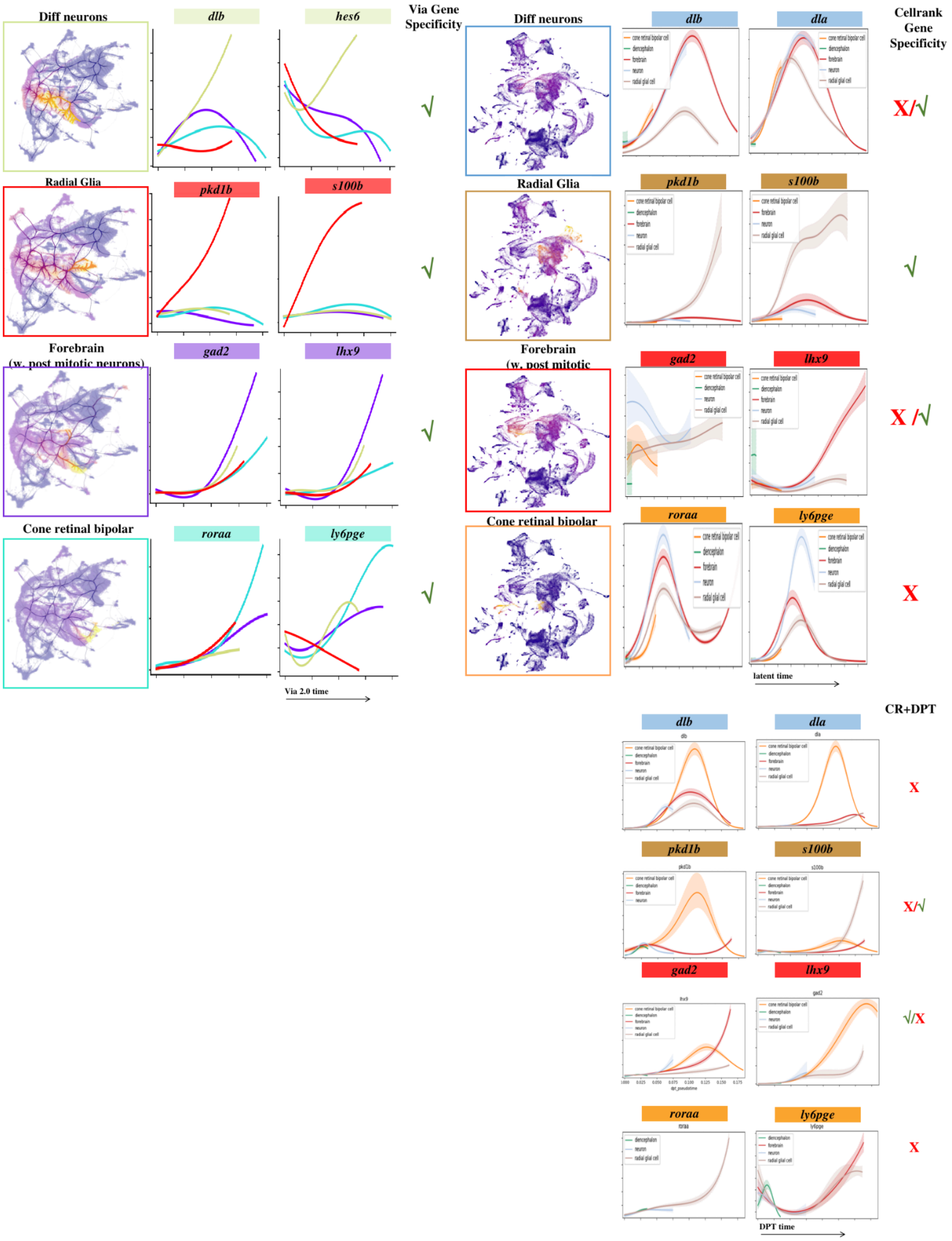


***Supplementary Fig.S8. Zebrahub neural ectoderm: Comparison of lineage paths and gene specificity trends.*** *Same as Fig.S7 but for the neural ectoderm lineages. Cell fates are manually assigned to CellRank in the case of forebrain and differentiation neurons.*

**Supplementary Fig S9. Zebrahub Non-neural Ectoderm lineage paths and gene trends**

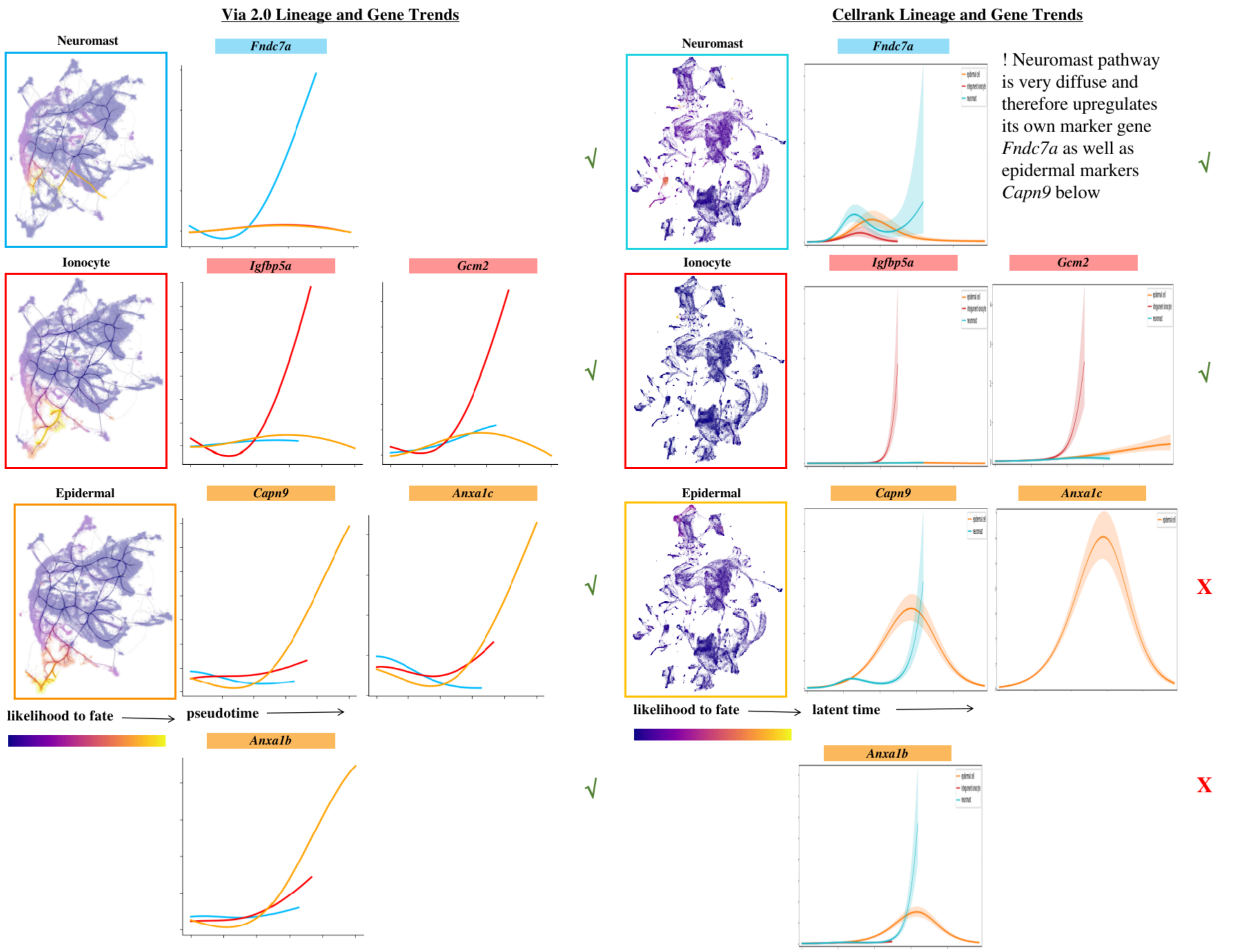


***Supplementary Fig.S9. Zebrahub non-neural ectoderm: Comparison of lineage paths and gene specificity trends.*** *Same as Fig.S8 but for the non-neural ectoderm lineages. Cell fates for Ionocyte and Neuromasts are manually assigned to Cellrank as they are not automatically detected.*

### Supplementary Fig S10. Impact of Memory on Mesodermal Lineage Probabilities (Zebrahub)

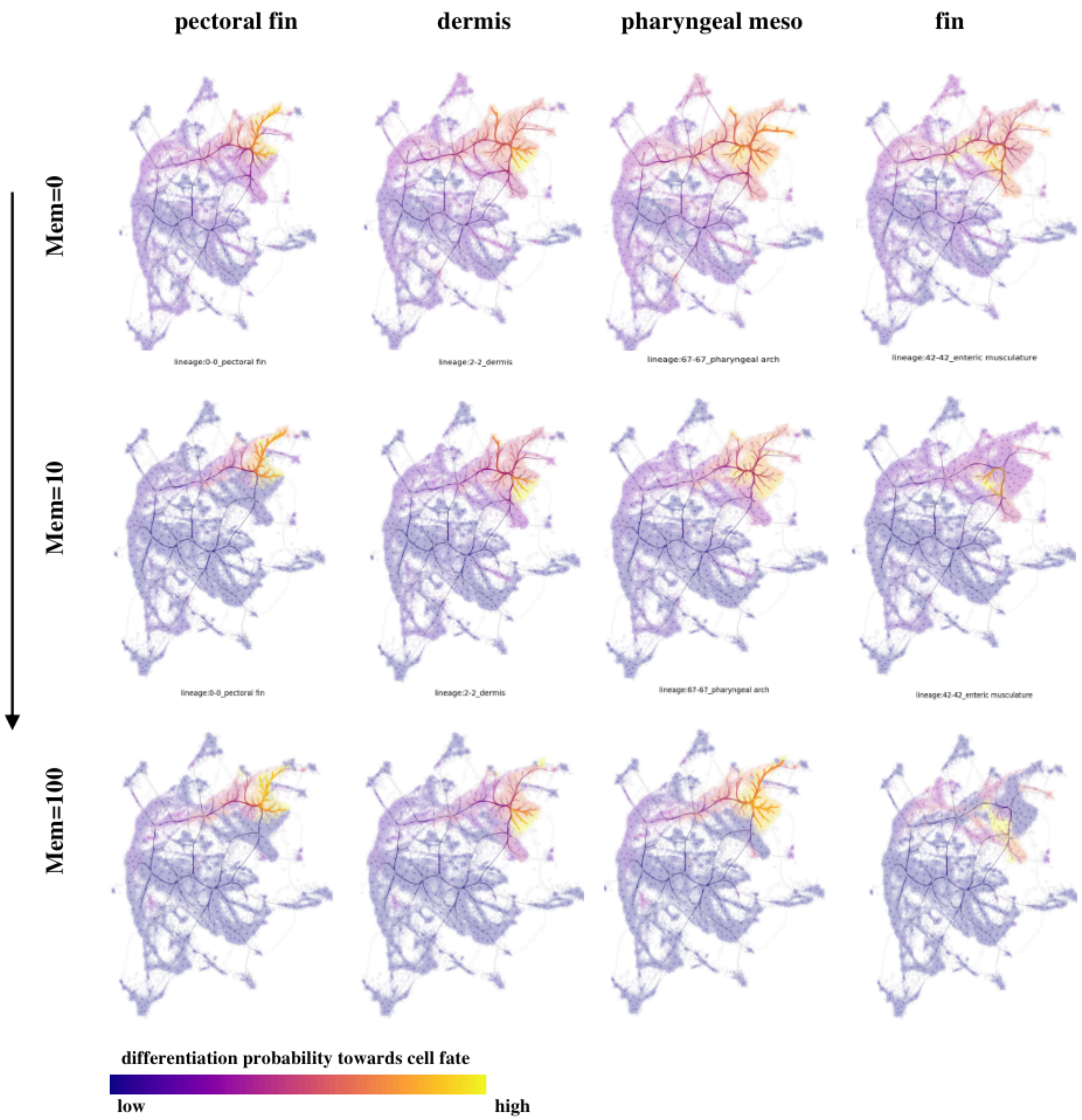


***Supplementary Fig S10. Impact of Memory on Mesodermal Lineage Probabilities:*** *When the random walk has no memory (is first order), we see that the pharyngeal and fin lineages have significant cell-fate probabilities in the non-mesodermal middle and lower branches, indicated by the light purple-pink coloration in these lower branches. As memory is increased the middle and lower ectoderm branch become a darker blue and the cells residing on them do not express a likelihood towards the mesodermal lineages. Including random walks with memory therefore helps refine the end-to-end pathways to cell fates*

**Supplementary Fig S11. Impact of memory on mesodermal gene trends (Zebrahub)**

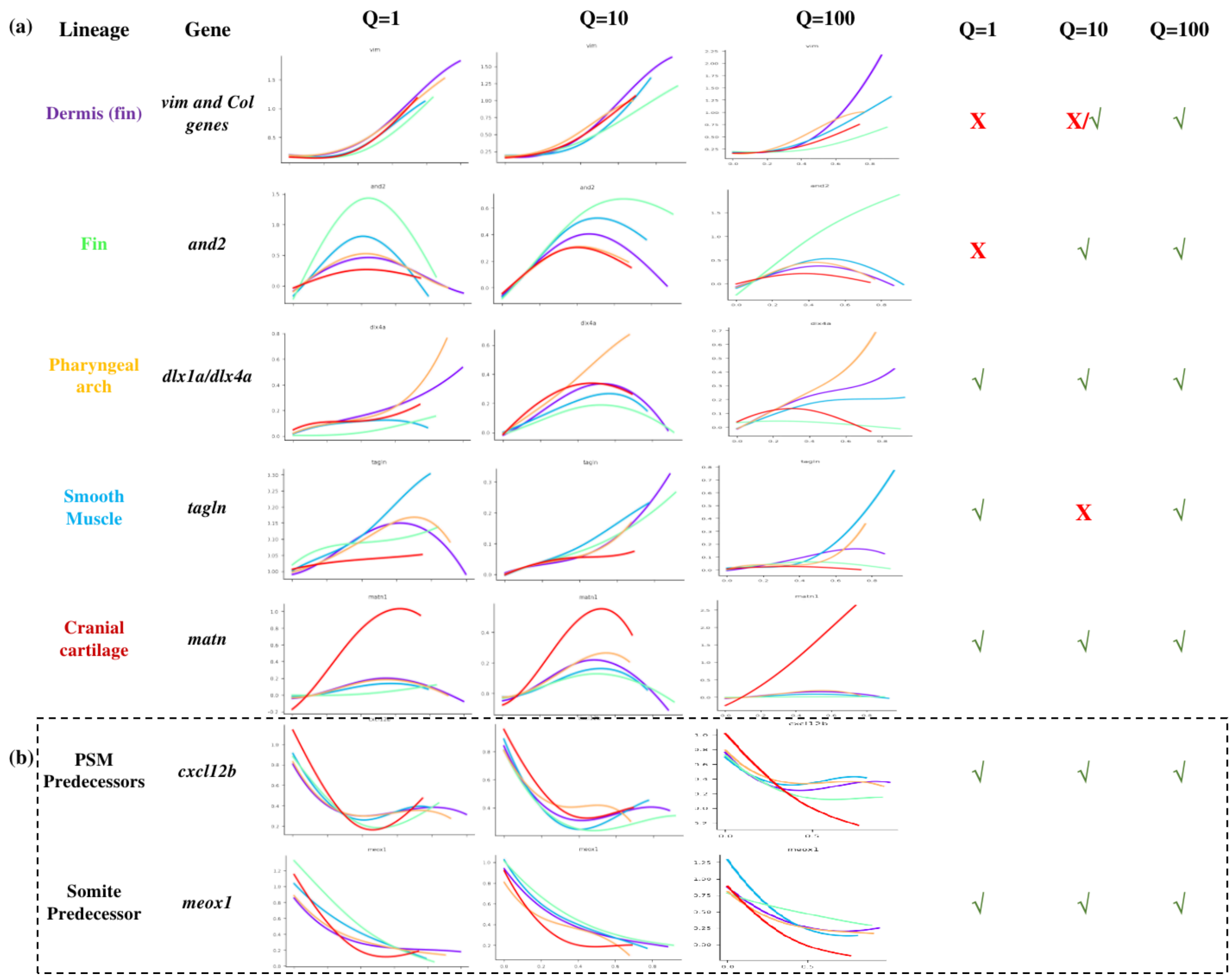


***Supplementary Fig S11. Impact of Memory on Mesodermal Gene Trends (Zebrahub):*** *(a) When the random walk has no memory (is first order), we see that the pharyngeal and fin lineages have significant cell-fate probabilities in the non-mesodermal middle and lower branches, indicated by the light purple-pink coloration in these lower branches. As memory is increased the middle and lower ectoderm branch become a darker blue and the cells residing on them do not express a likelihood towards the mesodermal lineages. Including random walks with memory therefore helps refine the end-to-end pathways to cell fates. (b) The predecessor genes are correctly shown to all be downregulated along the pseudotime axis.*

Fig.S12 Comparison of Visualization methods colored by stage

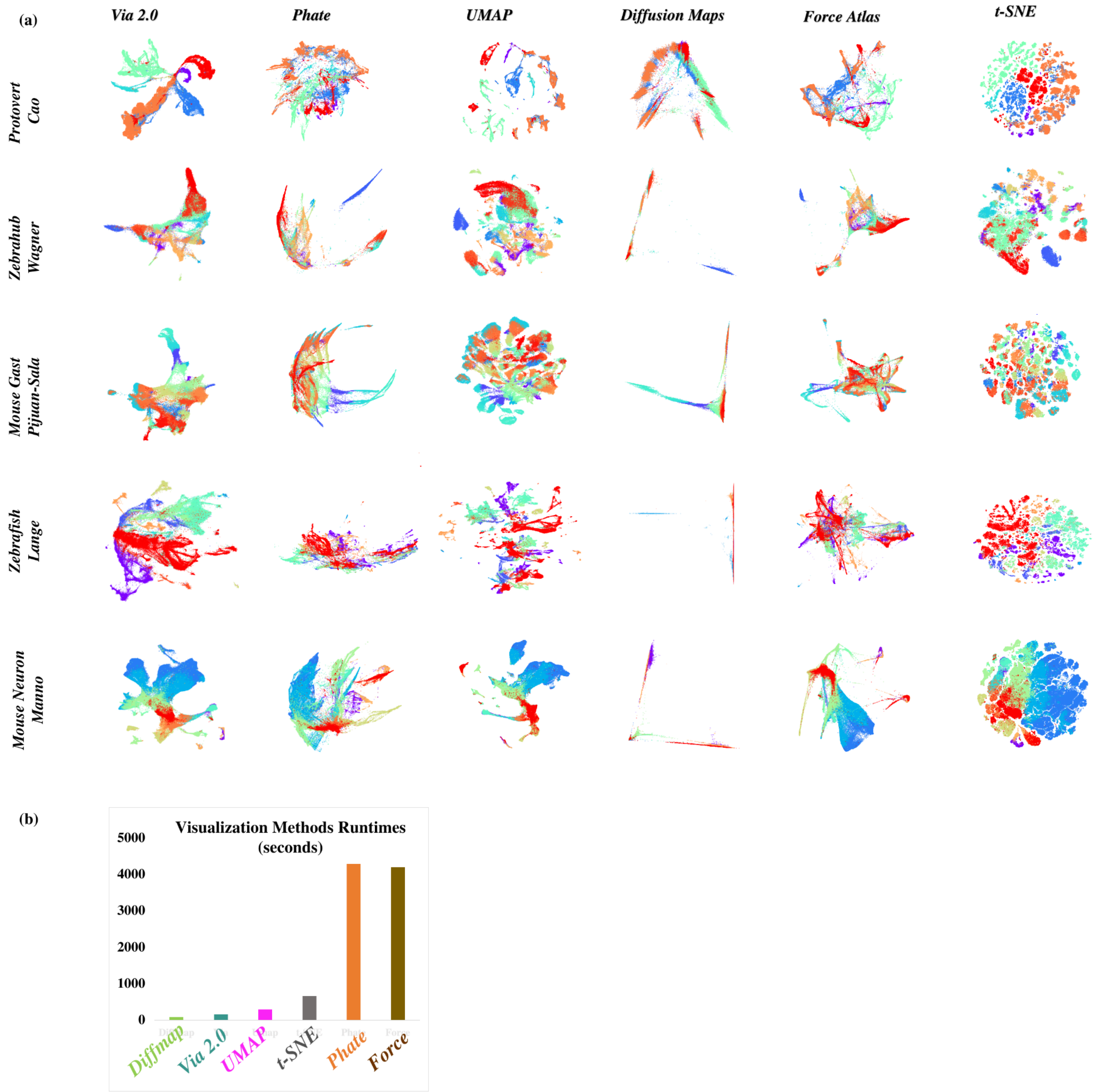


***Supplementary Fig.S12 Comparison of visualization methods on different time-series RNA-seq datasets.*** *(a) Colored by known tissue type. (b) Comparison of runtime in seconds to compute the single cell embeddings for Zebrahub (120K cells)*

**Fig.S13 Comparison of Visualization methods colored by stage**

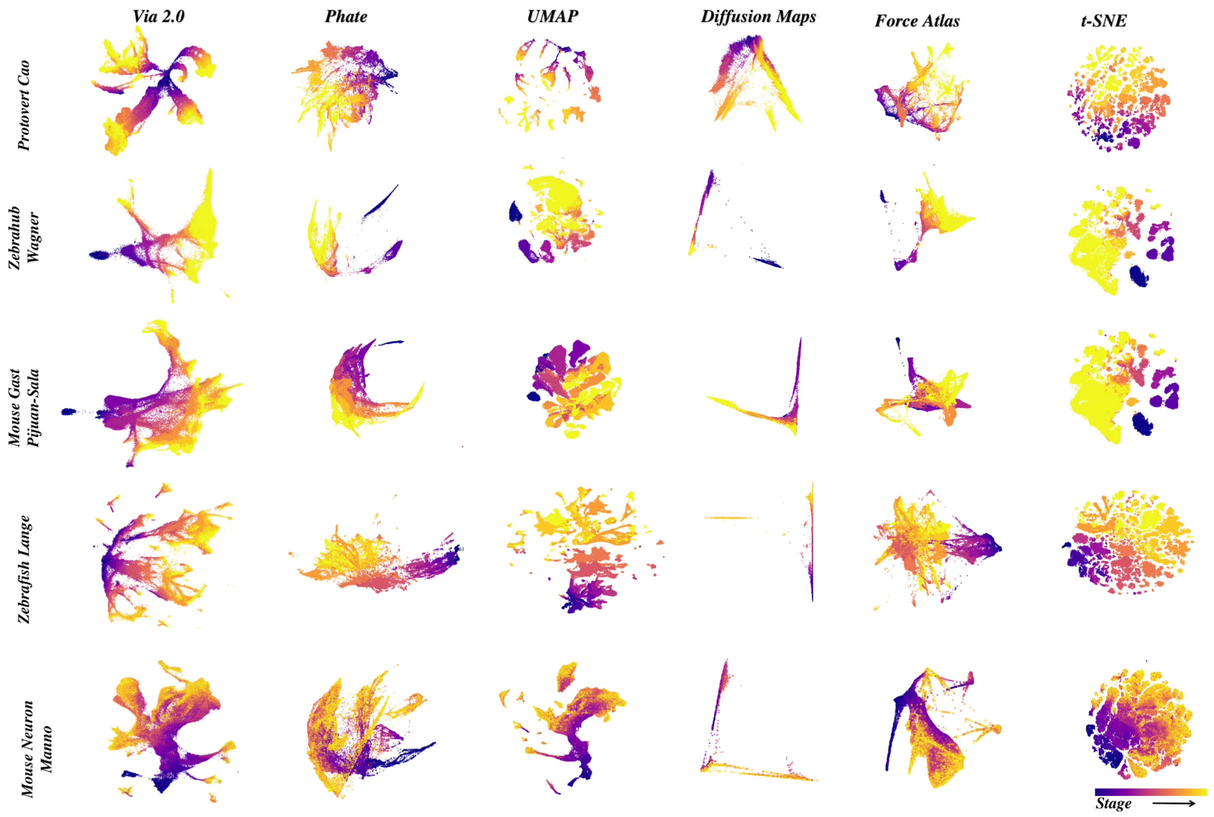


***Fig.S13 Comparison of visualization methods on different time-series RNA-seq datasets.*** *Colored by known developmental stage.*

**Fig.S14. Impact of key steps in Via2.0 Atlas embedding**

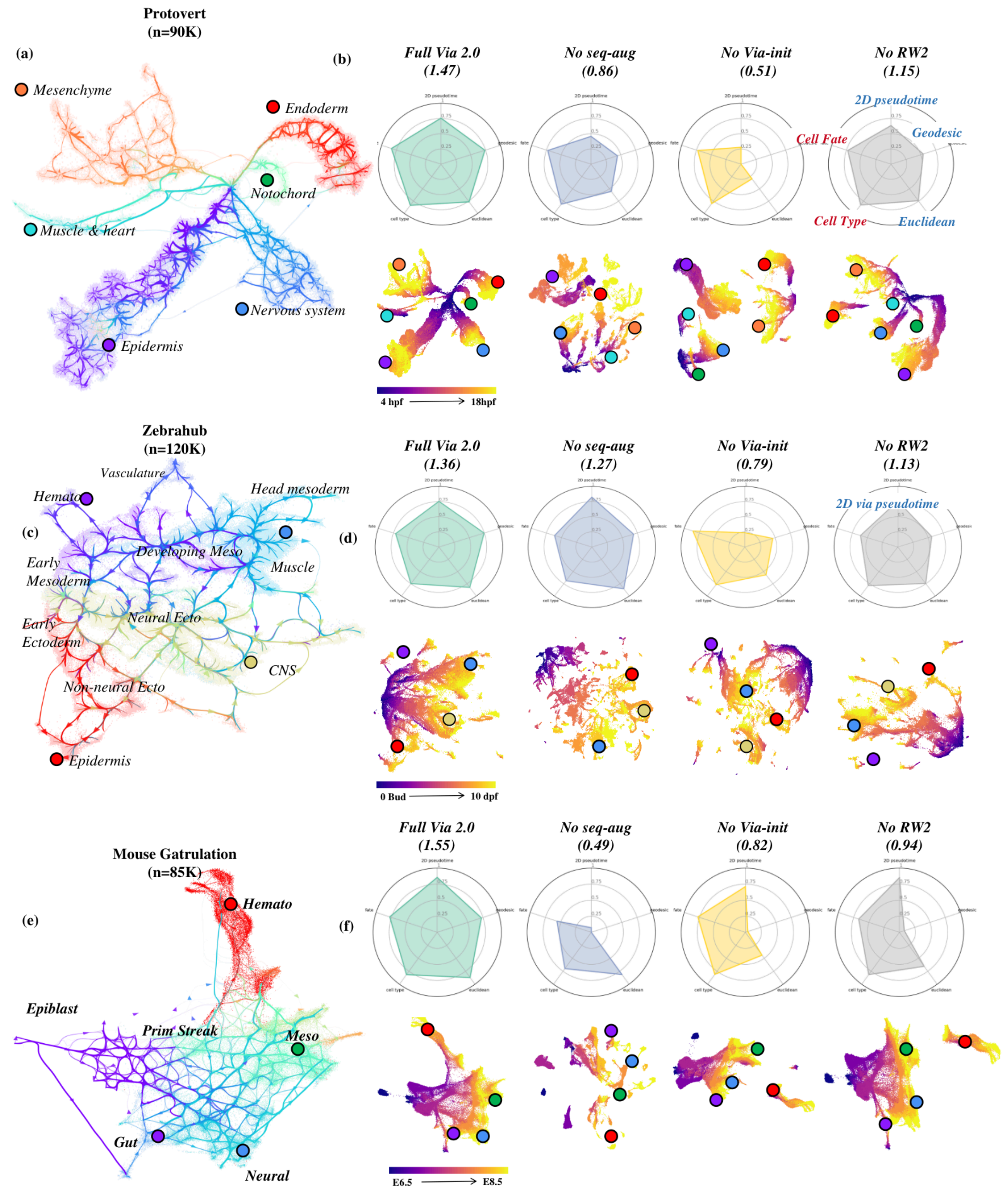


***Fig.S14. Impact of key steps in Via2.0 Atlas embedding*** *(a) Via 2.0 Atlas view colored by cell type for Ascidian embryo (b) showing the impact on quantitative score and quality of visual embedding colored by stage, when knocking-out one at a time, one of the main steps in generating the embedding. "Full" is the embedding score with all steps intact. "No-seq-aug" does not augment the sc-KNN graph with the known time-series information. "No Via-Initialization" skips the step where the embedding layout is otherwise initialized using the force-directed layout of the forward-biased TI cluster graph. "No-RW2" means that PCs are used instead of the same number of features from the graph-based node2Vec feature representation. (c-d) Zebrahub (e-f) Mouse Gastrulation*

### Fig.S15 Impact of Steps in Via 2.0 Embedding (stage)

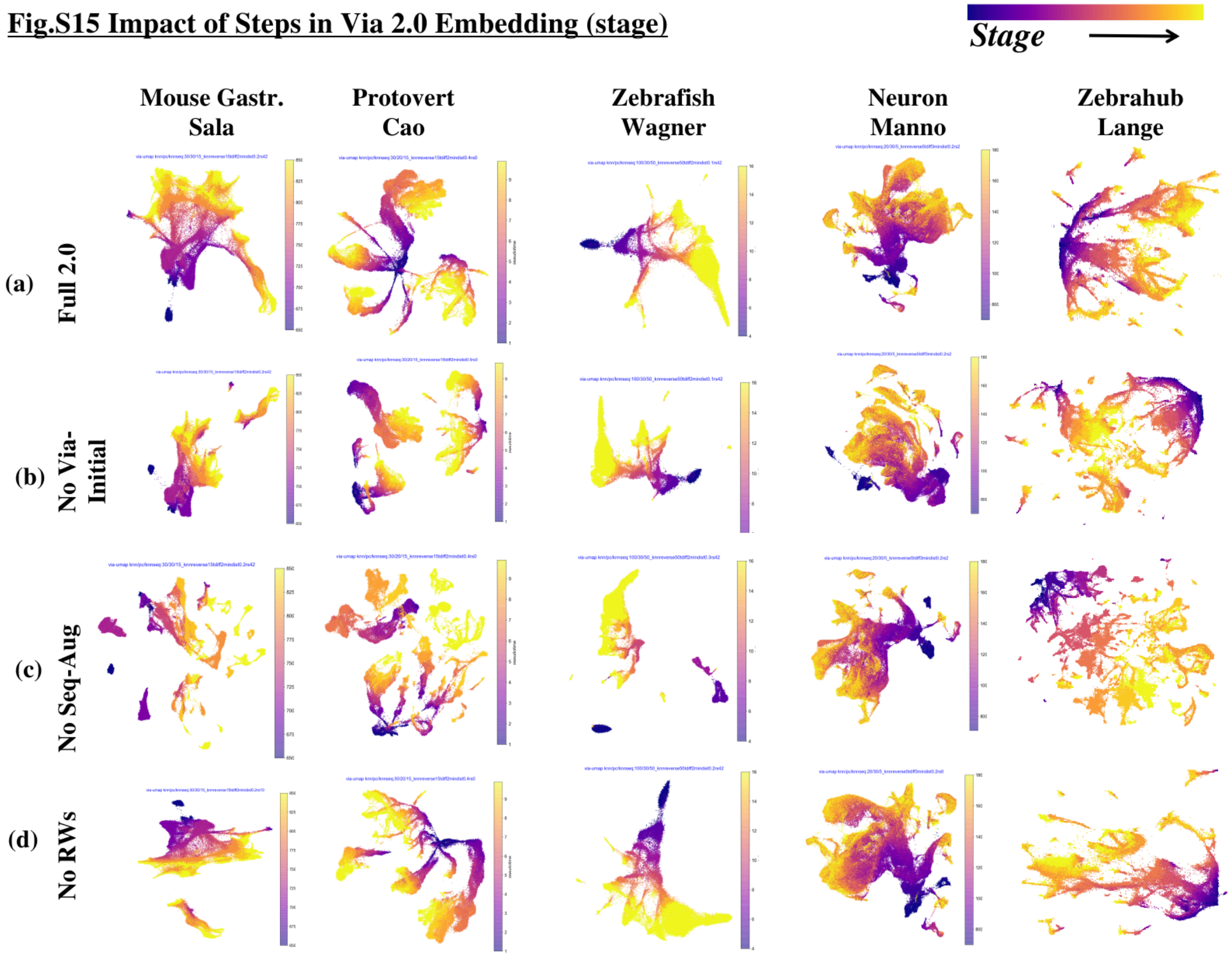


***Fig.S15 Impact of Steps in Via 2.0 Embedding (cells colored by known developmental stage)*** *Each row represents the effect of removing one of the key steps in the embedding computation. (a) Includes all key steps (b) Skips via-initialization of embedding using TI-directed and weighted cluster-graph layout (c) Skips leveraging experimental sequential labels to sequentially augment sc-KNN graph (d) uses PCs instead of Node2Vec feature representation (same number of PCs as RW2 features)*

**Supplementary Fig.S16. Impact of Steps in Via 2.0 Embedding (celltype)**

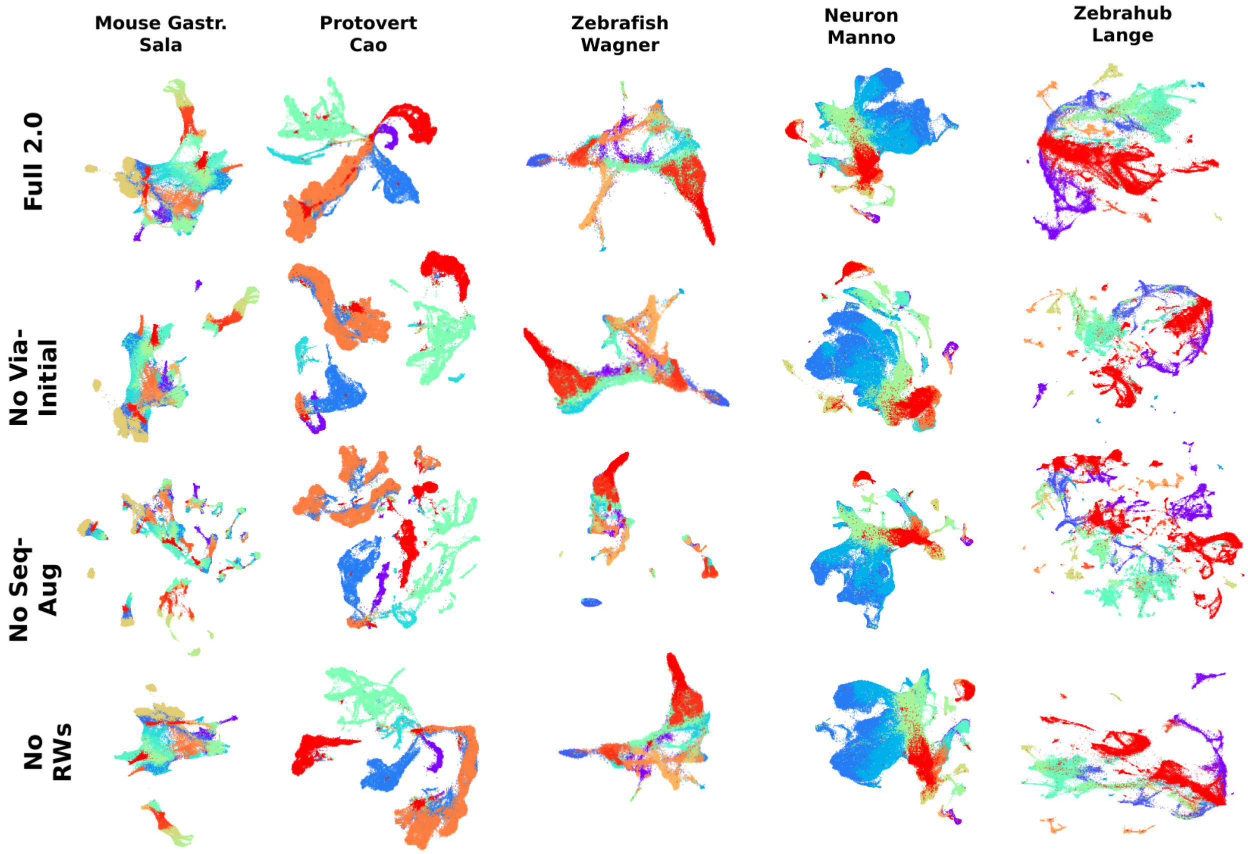


***Fig.S16 Impact of Steps in Via 2.0 Embedding (cells colored by tissue type)*** *Each row represents the effect of removing one of the key steps in the embedding computation. (a) Includes all key steps (b) Skips via-initialization of embedding using TI-directed and weighted cluster-graph layout (c) Skips leveraging experimental sequential labels to sequentially augment sc-KNN graph (d) uses PCs instead of Node2Vec feature representation (same number of PCs as RW2 features)*

### Fig.S17 Impact of Steps in Via 2.0 Embedding

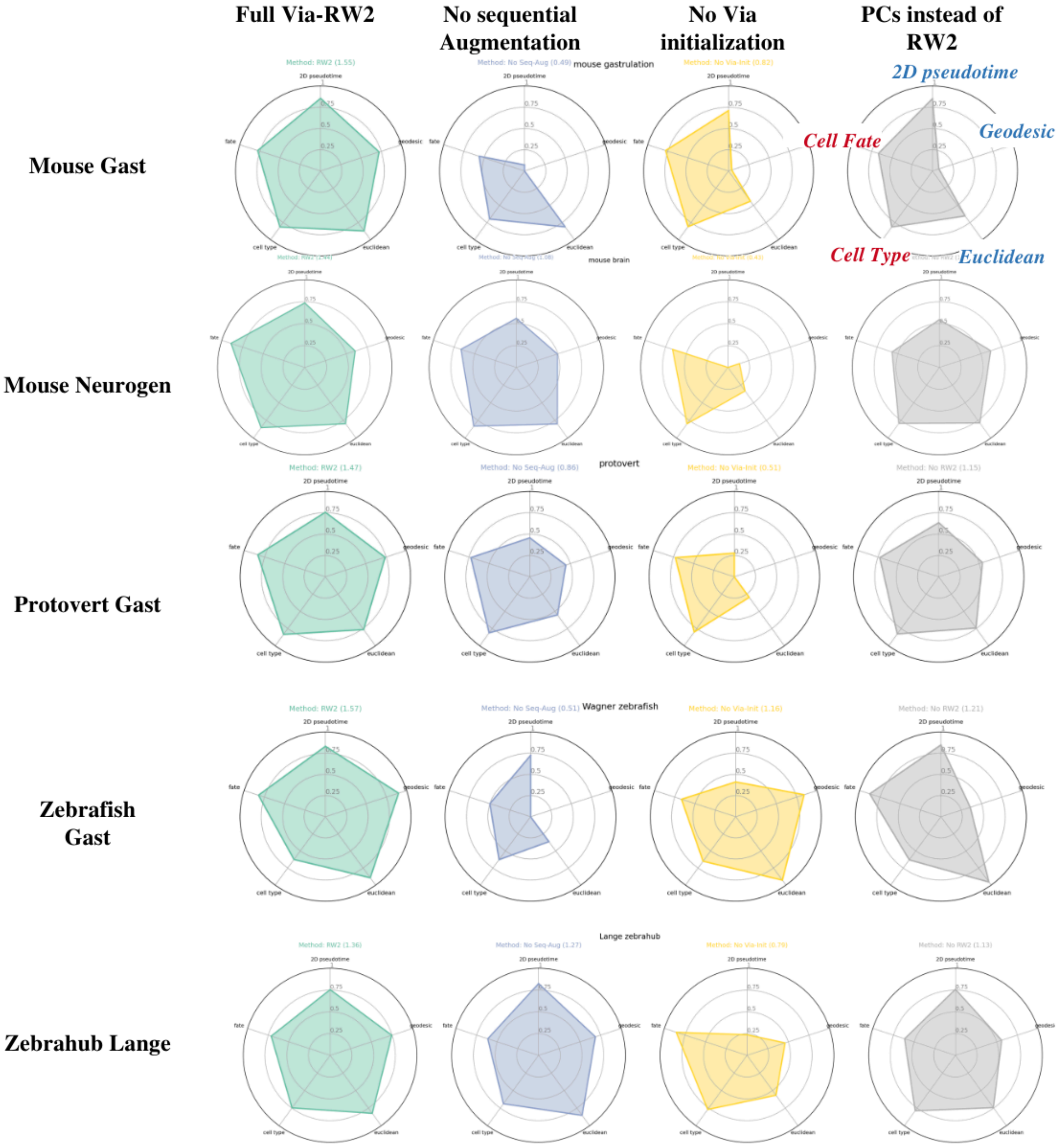


***Fig.S17 Impact of Steps in Via 2.0 Embedding on different time-series RNA-seq datasets.*** *Radar plots scoring preservation of structural continuity (blue metrics: 2D pseudotime, Geodesic root-cell correlation with developmental stage, Euclidean root-cell distance correlation with developmental stage) and cell type separation (red metrics: Cell Fate, Cell Type). Each column represents the effect of removing one of the key steps in the embedding computation. (Column 1) Includes all key steps (Column 2) Skips via-initialization of embedding using TI-directed and weighted cluster-graph layout (Column 3) Skips leveraging experimental sequential labels to sequentially augment sc-KNN graph (Column 4) uses PCs instead of Node2Vec feature representation (same number of PCs as RW2 features)*

**Supplementary Fig.S18 Second order random walk with memory**

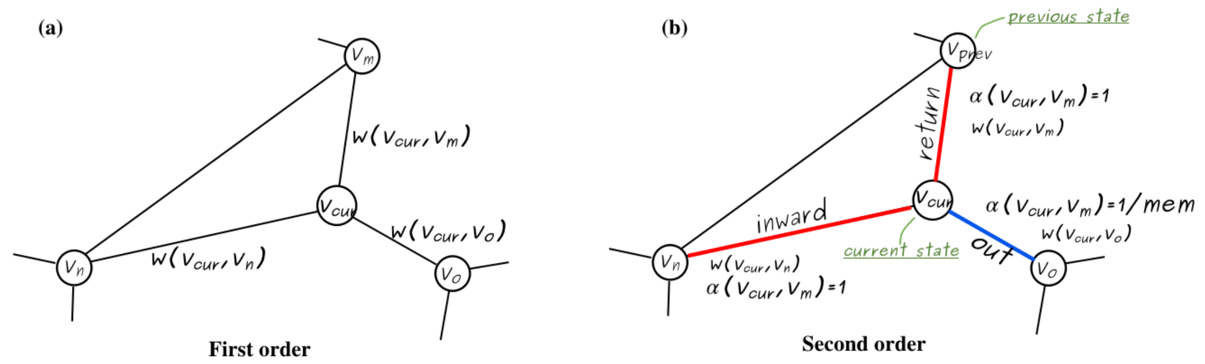


*Fig.S18 Second order random walk with memory*
